# Steroid hormone catabolites activate the pyrin inflammasome through a non-canonical mechanism

**DOI:** 10.1101/2021.10.29.466454

**Authors:** Flora Magnotti, Daria Chirita, Sarah Dalmon, Amandine Martin, Pauline Bronnec, Jeremy Sousa, Olivier Helynck, Wonyong Lee, Daniel Kastner, Jae Jin Chae, Michael F. McDermott, Alexandre Belot, Michel Popoff, Pascal Sève, Sophie Georgin-Lavialle, Hélène Munier-Lehmann, Tu Anh Tran, Ellen De Langhe, Carine Wouters, Yvan Jamilloux, Thomas Henry

## Abstract

The pyrin inflammasome acts as a guard of RhoA GTPases and is central to immune defences against RhoA-manipulating pathogens. Pyrin activation proceeds in two steps. Yet, the second step is still poorly understood. Using cells constitutively activated for the pyrin step 1, a chemical screen identified etiocholanolone and pregnanolone, two catabolites of testosterone and progesterone, acting at low concentrations as specific step-2 activators. High concentrations of these metabolites fully and rapidly activated pyrin, in a human-specific, B30.2 domain-dependent manner and without inhibiting RhoA. Mutations in *MEFV*, encoding pyrin, cause two distinct autoinflammatory diseases (PAAND and FMF). Monocytes from PAAND patients, and to a lower extent from FMF patients, displayed increased responses to these metabolites. This study provides a new perspective on pyrin activation, indicates that endogenous steroid catabolites can drive autoinflammation, through the pyrin inflammasome, and explains the “steroid fever” described in the late 1950s, upon steroid injection in humans.

## Introduction

Inflammasomes are innate immune complexes that contribute to antimicrobial responses ^1^ but can also be deleterious in various chronic autoinflammatory conditions ^2^. Inflammasome activation results in activation of the inflammatory caspase-1, cleavage of the pore forming protein GasderminD (GSDMD) that triggers a fast cell death, termed pyroptosis, and release of the inflammatory cytokines IL-1β and IL-18. Inflammasome sensors can act as direct pathogen-associated molecular pattern (PAMPs) receptors but have also evolved more general sensing mechanisms allowing them to be activated in response to damage-associated molecular patterns (DAMPs) or to homeostasis-altering molecular processes (HAMPs) ^3^. Furthermore, the NLRP3 inflammasome is regulated by several metabolites ^4^. Whether metabolites regulate other inflammasome sensors is currently unknown.

Pyrin is an inflammasome sensor acting as a guard of Rho GTPases activity ^5^. Bacterial toxins (e.g. *Clostridioides difficile* toxins A and B (TcdA/B)) or bacterial effectors (e.g. *Yersinia* YopE/T ^6, 7^) inhibit RhoA and trigger pyrin inflammasome activation. Similarly, by disrupting RhoA prenylation, mevalonate kinase deficiency activates pyrin ^8^. RhoA inhibition lifts the dynamic blockage of pyrin. Indeed, at steady state, pyrin is phosphorylated by PKN1/2 on two serine residues (S208 and S242), allowing a phospho-dependent chaperone from the 14-3-3 family to sequester pyrin away from downstream inflammasome molecules. PKN1/2, two kinases from the PKC superfamily, are RhoA effectors that are active in cells with homeostatic levels of RhoA activation. Inhibition of RhoA leads to loss of PKN1/2 activity, dephosphorylation of pyrin and ultimately triggers activation of the pyrin inflammasome ^5, 8–10^. Yet, pyrin dephosphorylation is not sufficient to trigger the full pyrin inflammasome activation ^11^.

We have therefore proposed a two-step model for pyrin activation (Fig. 1A). The first step corresponds to pyrin dephosphorylation, while the second step, which remains poorly understood, corresponds to the formation of ASC oligomers and caspase-1 activation. Interestingly, this model is consistent with the observation that colchicine and, other microtubule-depolymerising drugs, block pyrin activation downstream of its dephosphorylation ^10, 12^.

**Figure 1:**
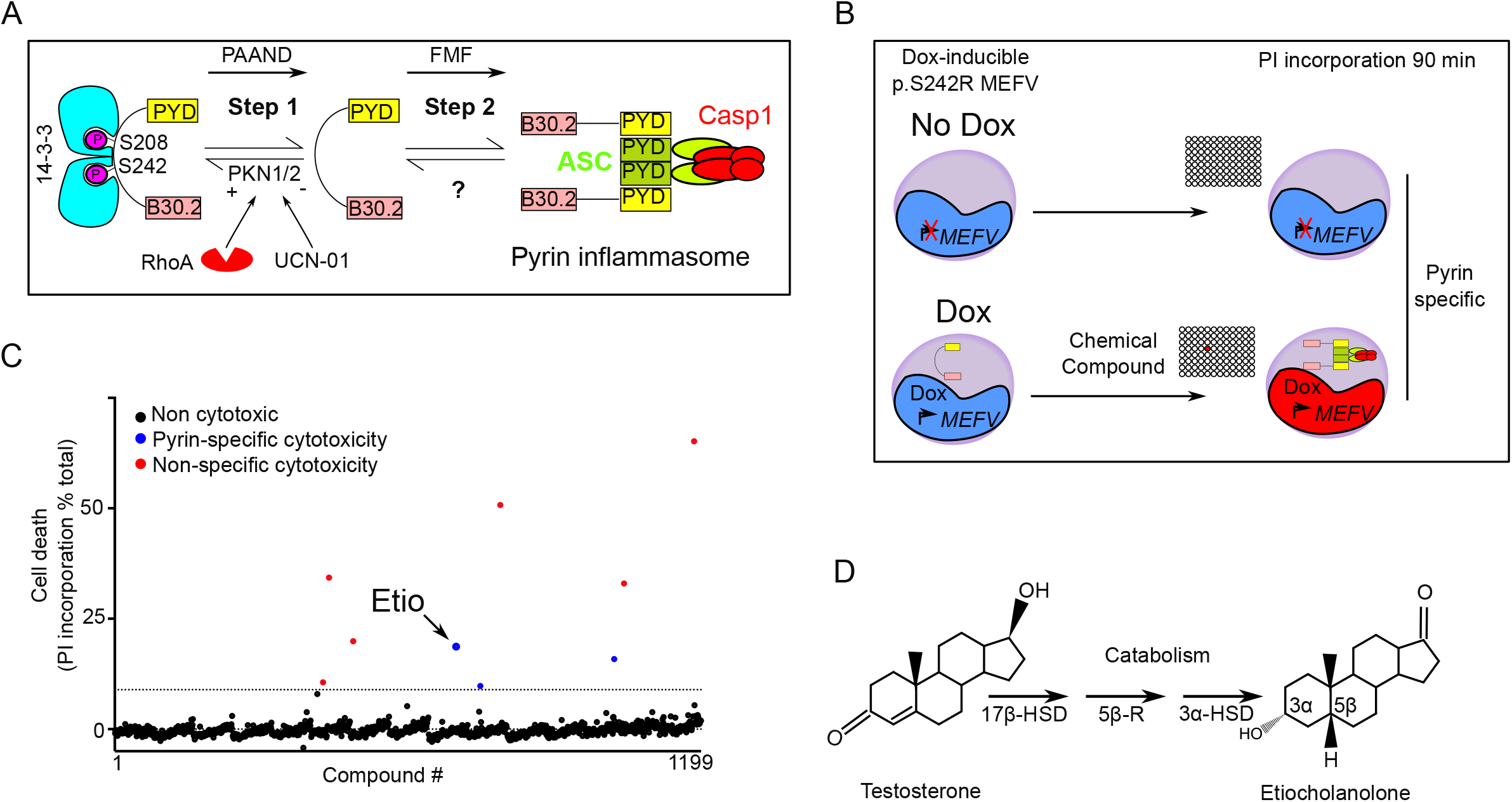
a chemical screen identifies etiocholanolone, a testosterone catabolite, as a pyrin inflammasome step 2 activator. (A) Model for pyrin two step activation mechanism. Step 1 is due to dephosphorylation of pyrin and loss of 14-3-3 binding and is constitutive in PAAND patient. Step 2 is uncharacterized but is upstream of ASC speck formation and is constitutive in FMF patients. (B) Chemical screen overview: cells expressing (Dox: doxycycline) or not (No Dox) p.S242R *MEFV* were exposed to individual chemical compounds. 90 min post- exposure, cell death was monitored using propidium iodide (PI). Compounds driving cell death independently of pyrin (i.e. in the absence of doxycycline) were excluded. (C) Screen results are shown, each dot represents the cell death value of cells exposed to one chemical compound. The dotted line represents the mean + 3 SD. Red dots represent non-specific hits (killing cells irrespective of the presence or the absence of Dox) while blue dots represent specific hits displaying cytotoxicity only upon pyrin expression. Etiocholanolone (Etio, 6.9 μM) is highlighted. (D) Structure of progesterone and its catabolite etiocholanolone are shown. The stereochemistry of carbon 3 and 5 is indicated. 17β-HSD: 17β-hydroxy-steroid dehydrogenase, 5β-R: 5β-reductase, 3α-HSD: 3α-hydroxy-steroid dehydrogenase.

The two-step model is also supported by the two distinct autoinflammatory diseases associated with mutations in *MEFV*, the gene encoding pyrin ^13^. Indeed, mutations in *MEFV* exon 10, coding the B30.2 domain of pyrin, cause Familial Mediterranean Fever (FMF) and are associated with a constitutively activated step 2 ^11^. In contrast, mutations affecting the serine residues, which are dephosphorylated during step 1 ^9, 14^, or the neighbouring residues required for the interaction with 14-3-3 proteins ^15^ cause Pyrin-Associated Autoinflammation with Neutrophilic Dermatosis (PAAND).

Despite its importance in health and disease, the mechanism responsible for step 2 is still largely unclear and this study aimed at increasing our knowledge on this specific regulatory mechanism.

## Results

### A chemical screen identified sex hormone catabolites as pyrin step 2 activators

To identify molecules that trigger step 2 of the pyrin inflammasome cascade, we used a human monocytic cell line (U937) expressing the PAAND *MEFV* variant (i.e. p.S242R) under the control of a doxycycline-inducible promoter ^11^ (Fig. 1B). In these cells, the p.S242R mutation mimics pyrin dephosphorylation resulting in a pyrin protein constitutively activated for step 1. Cells expressing (in the presence of doxycycline) p.S242R *MEFV* were treated with compounds from the Prestwick library (n=1199). At 90 minutes post-addition, cell death was measured by quantifying propidium iodide incorporation. A counter-screen was performed simultaneously in the same conditions but in the absence of doxycycline, (i.e. in the absence of pyrin expression) to retain compounds triggering pyrin-specific cell death. One compound, etiocholanolone (6.9 μM) was identified as triggering a fast cell death in a doxycycline-dependent manner (Fig. 1C). Etiocholanolone, also termed 3α-hydroxy 5β-androstan-17-one, is an endogenous catabolite of the steroid hormone, testosterone (Fig. 1D). Except for medrysone, which was weakly active, none of the other steroids in the Prestwick library triggered pyrin-specific cell death.

The ability of etiocholanolone to trigger cell death in a p.S242R-*MEFV*-dependent manner was validated in an independent set of experiments and demonstrated to be dose-dependent (Fig. 2A). To assess the specificity of etiocholanolone and perform structure-activity relationship studies, a number of hormones and catabolites were then tested. Interestingly, pregnanolone (3α-hydroxy-5β-pregnan-20-one), a catabolite of progesterone, triggered p.S242R-*MEFV*-dependent cell death at an even lower concentration than etiocholanolone (Fig. 2A). In contrast, neither testosterone nor progesterone triggered cell death, even at high concentrations (half maximal effective concentration EC50>1000 μM). Furthermore, both cortisol and its catabolite, tetrahydro-cortisol, were inactive (Fig. 2B), demonstrating the specificity of the pregnanolone and etiocholanolone in triggering this response.

**Figure 2:**
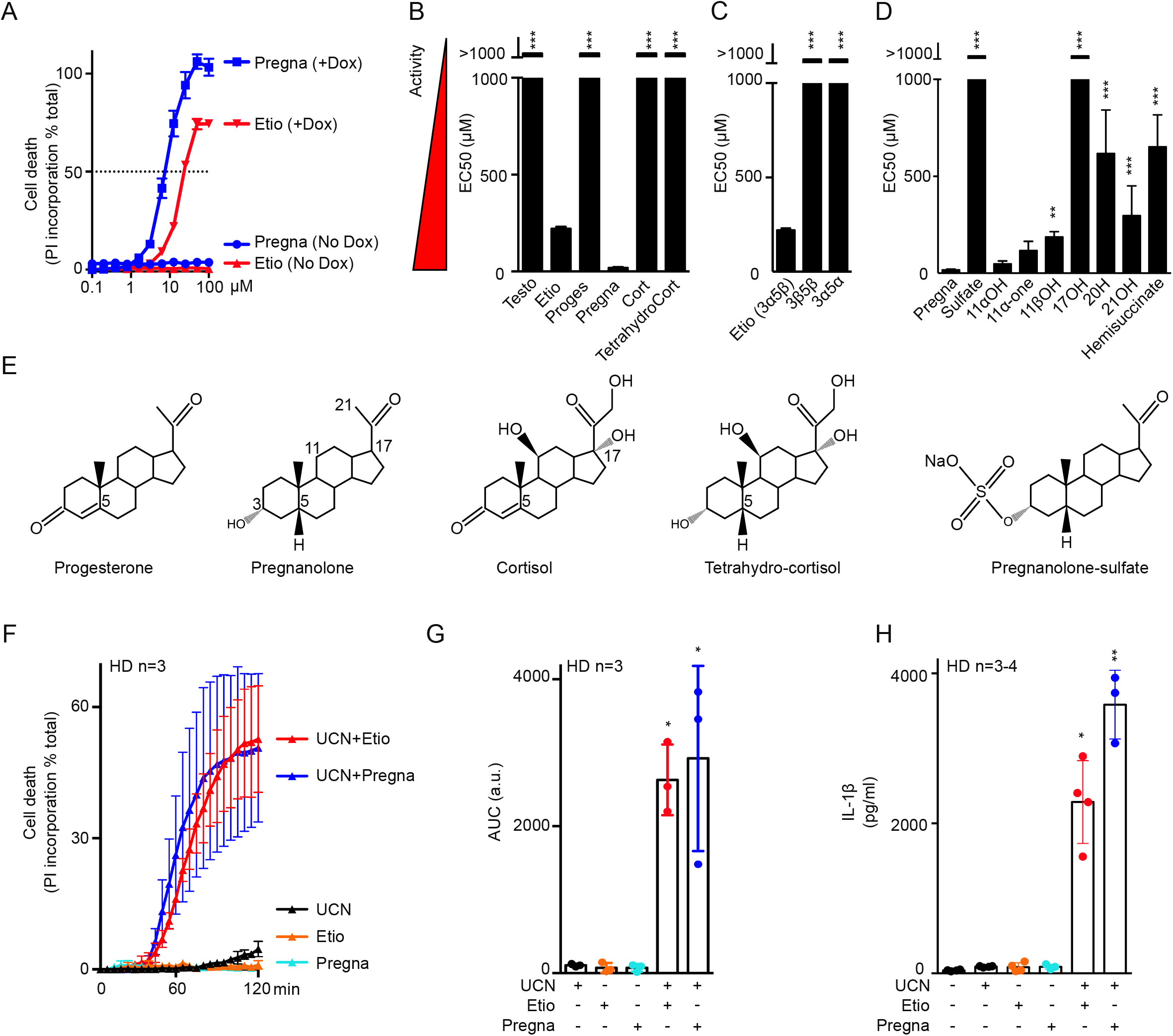
Pregnanolone and etiocholanolone specifically triggers pyrin inflammasome step 2. (A) Pregnanolone (Pregna, blue) and etiocholanolone (Etio, red) were added at different concentrations on cells expressing (in the presence of doxycycline, DOX+) or not (No Dox) p.S242R *MEFV*. Cell death was determined at 3 h post-addition. The concentration triggering 50% cell death (horizontal dotted line) determined the EC50 (Half maximal effective concentration which is inversely correlated to the activity of the tested molecule). (B) Structure activity of various steroid hormones, their catabolites, (C) of etiocholanolone (also known as 3α-hydroxy 5β-androstan-17-one (3α, 5β)) and its two stereoisomers 3β-hydroxy-5β-androstan-17-one (3β, 5β) and androsterone (3α, 5α), (D) of pregnanolone and molecules closely related. EC50 were calculated as in A. (E) Structures of selected compounds. All compounds can be found in supplemental Fig. S1. (F) Primary monocytes from healthy donors (HD, n=3) were pre-treated with pregnanolone (6 μM) or etiocholanolone (12 μM) for 1 h followed by addition of the PKC superfamily inhibitor, UCN-01. Cell death was monitored every 5 min for 2 h. (G) The Area Under the Curve (AUC) was computed for each HD. (H) Primary monocytes from healthy donors (HD, n=3-4) were treated with LPS for 2 h, then pre-treated with pregnanolone (6 μM) or etiocholanolone (12 μM) for 1 h followed by UCN-01 addition. IL-1β concentration in the supernatant was quantified at 3 h post-UCN-01 addition. <u>Data information:</u> One experiment representative of three (A) or two (B-D) independent experiment is shown. Mean and SEM of triplicates are shown. Testo: testosterone; Proges: progesterone; Cort: cortisol; TetrahydroCort: tetrahydro-cortisol; Sulphate: 5β-pregnan-3α-ol-20-one sulphate; 11αOH: 5β-pregnan- 3α, 11α-diol-20-one; 11-one: 5-beta-pregnan-3α-ol-11, 20-dione; 11βOH: 5-β-pregnan-3-α, 11β-diol-20- one; 17OH: 5-β-pregnan-3-α, 17 diol-20-one; 20H: 5-β-pregnan-3-α-ol; 21OH: 5-β-pregnan-3-α, 21-diol- 20-one; Hemisuccinate: 5-β-pregnan-3-α, 21-diol-20-one 21 hemisuccinate. (B, C, D) One-way ANOVA with Dunn’s correction was applied. ***:p<0.001; **p=0.007. (F) each point corresponds to the mean +/- SEM of 3 HD values, each one being the mean of a triplicate. (G) Each point corresponds to the mean AUC of kinetics of one HD performed in triplicate, the bar represents the mean +/- SEM of 3HD. AUC are expressed as arbitrary units (a.u.). Friedman test with Dunn’s correction for multiple analysis was performed in comparison to untreated cells. *: p=0.023 (Etio + UCN); p=0.011 (Pregna + UCN). (H) Each point corresponds to the mean IL-1β concentration of 1 HD calculated from a triplicate, the bar represents the mean +/- SEM of 3-4 HD. Kruskal-Wallis test with Dunn’s correction for multiple analysis was performed in comparison to LPS-treated cells. *:p=0.011; **p=0.002.

Etiocholanolone and pregnanolone share the same stereochemistry on carbon 3 (3α) and 5 (5β) of the sterol. To evaluate the stereospecificity of the response, the two stereoisomers of etiocholanolone were tested for their ability to trigger p.S242R-*MEFV*-dependent cell death. 3β-hydroxy-5β-androstan-17-one (3β5β) and androsterone, (i.e. 3α-hydroxy 5α-androstan-17-one, (3α5α)), displayed EC50 greater than 1000 μM (Fig. 2C and supplemental Fig. S1), indicating that the response is stereospecific.

Furthermore, experiments with pregnanolone derivatives demonstrated that sulfation, a modification associated with steroid inactivation and excretion ^16^ leads to a complete loss of activity (Fig. 2D). All the modifications tested on the sterol ring or the terminal carbon or ketone (supplemental Fig. S1) decreased the pyrin-specific cytotoxicity. In particular, hydroxylation on C17 fully abolished the cell death, thereby explaining the absence of activity of the cortisol catabolite (Fig. 2D, E). Overall, these structure-function analyses demonstrated that etiocholanolone and pregnanolone are both highly specific compounds that trigger S242R-pyrin-mediated cell death.

To validate this result in primary human monocytes, we used the PKC superfamily inhibitor, UCN-01, which inhibits PKN1/2 and dephosphorylates pyrin, thus recapitulating the effect of the p.S242R mutation. As previously described ^11, 17^, UCN-01 alone does not trigger pyroptosis in primary human monocytes from healthy donors. Similarly, neither etiocholanolone (12 μM) nor pregnanolone (6 μM) alone were cytotoxic. Yet, the combination of UCN-01 and either of these two steroid catabolites triggered a very fast cell death (Fig. 2F, G) and IL-1β release (Fig. 2H). These results thus strongly suggest that, in primary monocytes, steroid catabolites activate pyrin step 2 and trigger activation of the pyrin inflammasome and pyroptosis, in the presence of the step 1 activator, UCN-01.

### High concentrations of steroid catabolites trigger full activation of the pyrin inflammasome in the absence of a step 1 activator

Interestingly, while at low concentration (<12 μM) steroid catabolites required activation of step 1, through either genetic (p.S242R) or chemical (UCN-01) means, we noticed that, at high concentrations, both etiocholanolone (Fig. 3A) and pregnanolone (Fig. 3B) triggered death of U937 cells expressing WT *MEFV*. High doses of pregnanolone (50 μM) or etiocholanolone (100 μM) triggered pyrin dephosphorylation to a similar extent as TcdA or UCN-01 while low doses of steroid catabolites did not (Fig. 3C). These results suggested that, at high doses, these molecules can trigger both step 1 and step 2. Accordingly, primary human monocytes exposed to high doses of steroid catabolites (50-100 μM) underwent a fast cell death (Fig. 3D, E) and released IL-1β (Fig. 3F). Primary neutrophils, treated with steroid catabolites, also underwent a rapid cell death (supplemental Fig. S2A). Etiocholanolone or pregnanolone addition triggered processing and release of caspase-1, IL-1β and GSDMD (Fig. 3G). Furthermore, pre-treatment with the caspase-1 inhibitor, VX765, fully abolished IL-1β release by primary human monocytes (supplemental Fig. S2B). Finally, Casp1^KO^ and GSDMD^KO^ U937 expressing WT pyrin were fully resistant to steroid catabolites-mediated cell death (Fig. 3H) and did not release substantial amounts of IL-1β (Fig. 3I). Altogether, these experiments demonstrated that steroid catabolites specifically triggered pyrin inflammasome activation and caspase-1-dependent, GSDMD-dependent pyroptosis and IL-1β release. This response was independent of NLRP3 (supplemental Fig. S2C).

**Figure 3:**
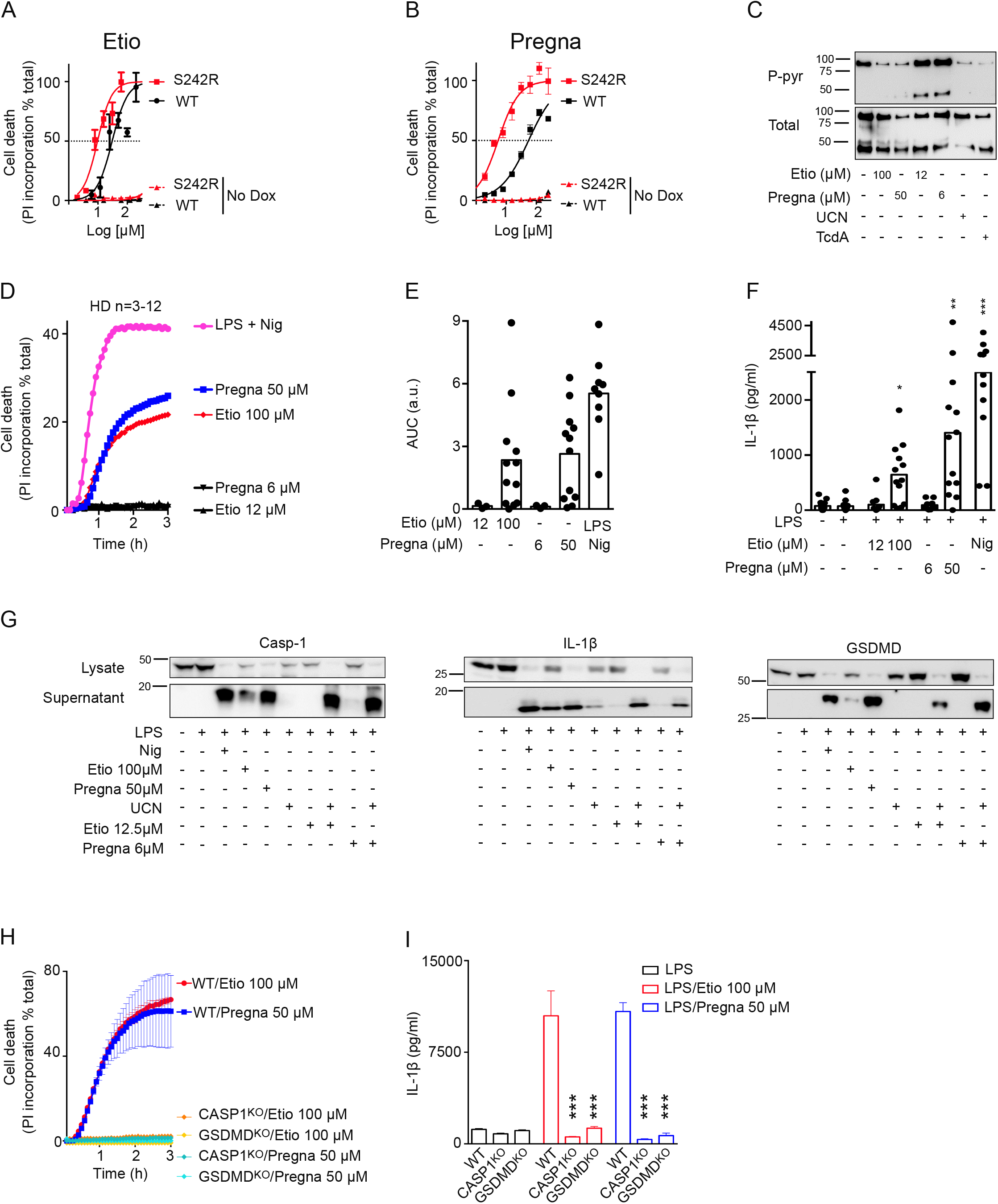
High concentrations of etiocholanolone and pregnanolone trigger full activation of pyrin inflammasome. (A, B) U937 cells expressing (plain lines) or not (No Dox, dotted lines) p.S242R (red) or WT (black) *MEFV* were treated with various concentration of etiocholanolone (A) or pregnanolone (B). Cell death was determined at 3 h post-addition. (C) 3xFlag-WT pyrin from U937 cells treated with the indicated stimuli at the indicated concentrations was immunoprecipitated. Ser242 phosphorylation (P-pyr) and total pyrin levels were monitored by Western blot. (D) Monocytes from healthy donors (n=3-12) were treated with low (black) or high concentrations of etiocholanolone (red) and pregnanolone (blue) or with LPS + Nigericin (Nig, Magenta). Propidium iodide incorporation (PI) was monitored every 5 min for 3 h. (E) The corresponding Area Under the Curve (AUC) for each donor are shown. (F) Monocytes were primed with LPS for 3 h and exposed to the indicated stimuli at the indicated concentration. IL-1β concentration in the supernatant was quantified at 3 h post-addition. (G) Monocytes from one HD were primed or not with LPS for 3 h before addition of the indicated stimuli. Caspase-1, IL-1β and GSDMD processing were analysed by Western blot in the cell lysate and supernatant at 3 h post stimuli addition. (H-I) *MEFV*-expressing U937 monocytes (H) or PMA-differentiated U937 macrophages (I) WT or knock-out for *CASP1* or *GSDMD* as indicated were treated with doxycycline during 16 h. (H) Propidium iodide (PI) incorporation was monitored every 5 min for 3 h post stimuli addition. (I) cells were primed with LPS for 3 h before addition of the indicated stimuli. IL-1β concentration in the supernatant was quantified at 3 h post-addition. <u>Data information:</u> One experiment representative of three (A-C) to two (H-I) independent experiment is shown. Mean and SEM of triplicates are shown. (A-B) Non-linear regression curve computed using least squares fit method is shown. (F) Kruskal-Wallis test with Dunn’s multiple comparisons tests was performed to compare the different treatments to the LPS treatment. *: p=0.026; **: p=0.0011; ***: p<0.001. (G) One-way ANOVA analysis with Sidak’s multiple comparisons test was performed to compared WT U937 to *CASP1*^KO^ or *GSDMD*^KO^ cells. ***:p<0.001.

### Steroid catabolites differ from the prototypical activator TcdA and trigger pyrin activation in a B30.2-dependent manner and in the absence of RhoA inhibition

Activation of the pyrin inflammasome in response to RhoA-inhibiting toxins depends on microtubule network integrity and is inhibited by microtubule-depolymerising drugs (e.g. colchicine or nocodazole) ^10, 12^. Similarly, colchicine and nocodazole inhibited pyroptosis induced by steroid catabolites in primary human monocytes (Fig. 4A, B). Colchicine also reduced IL-1β release in response to etiocholanolone and pregnanolone (Fig. 4C) while it had no detectable action on NLRP3-dependent pyroptosis.

**Figure 4:**
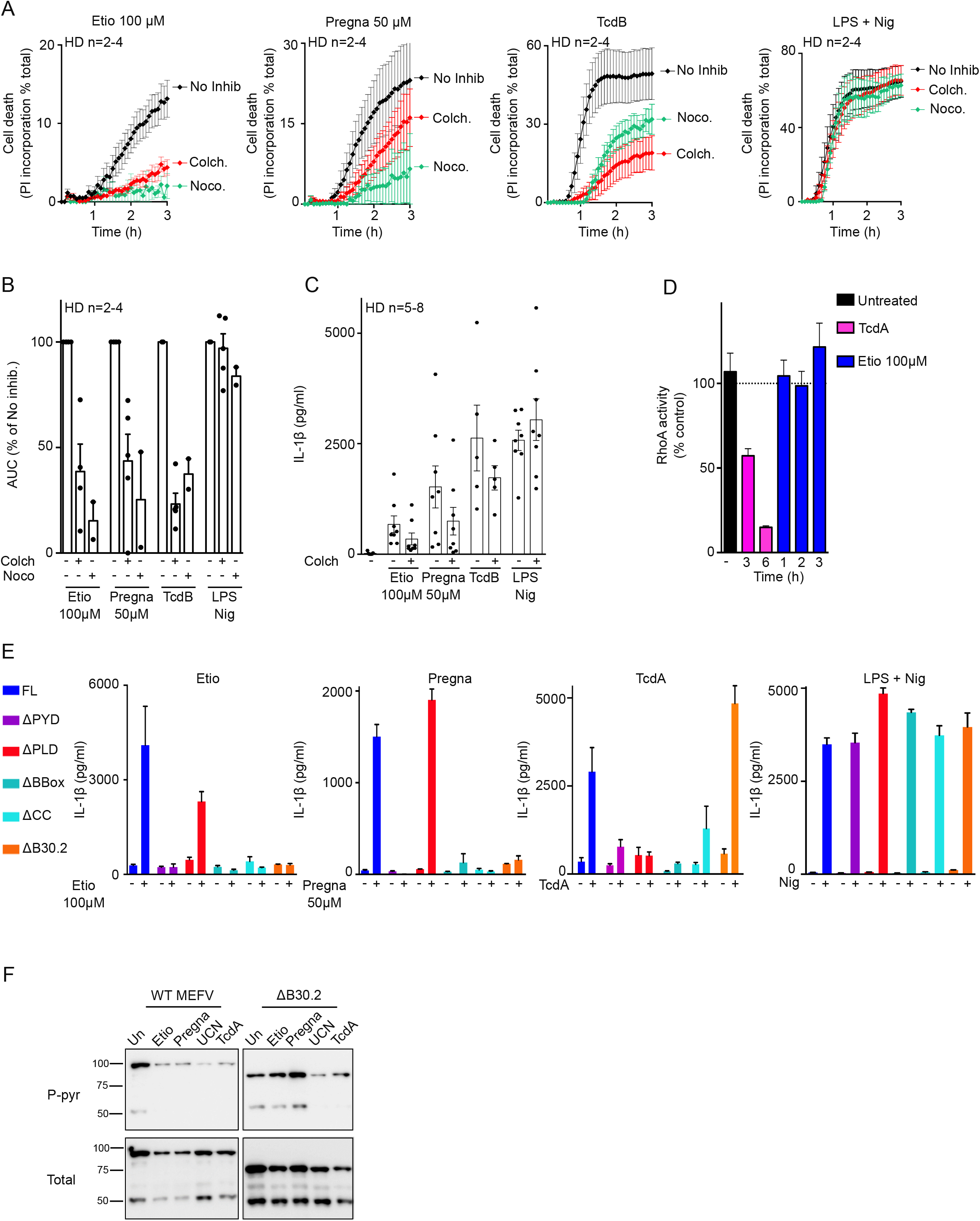
Pyrin inflammasome activation proceeds differently following TcdA/B and steroid catabolites addition. (A) Monocytes from HD (n=2-4) were treated with colchicine (Colch.) or nocodazole (Noco.) and 30 min later with the indicated stimuli. Propidium iodide incorporation was monitored every 5 min for 3 h. (B) The Area Under the Curve (AUC) are shown. For each HD, the AUC values in the presence of inhibitors were normalized to the AUC value obtained in the absence of inhibitor. (C) Monocytes from HD (n=5-8) were treated with colchicine (Colch.) and 30 min later with the indicated stimuli. IL-1β concentration in the supernatant was quantified at 3 h post-addition. (D) RhoA activity was determined by G-LISA at different time post-treatment in the lysate of U937 cells. The activity of the different treatments at the indicated time is presented in supplemental Fig. S3A. (E) Doxycycline-induced, PMA-differentiated U937 macrophages expressing *MEFV* Full- length (FL), deleted of the pyrin (ΔPYD), of the phosphorylated linker (ΔPLD), of the BBox (ΔBbox), of the Coiled-coil (ΔCC) or the B30.2 (ΔB30.2) domains were treated LPS for 3 h and then with the indicated stimuli. IL-1β concentration in the supernatant was quantified at 3 h post-addition. (F) Doxycycline-induced, U937 monocytes expressing WT or ΔB30.2 *MEFV* were treated with the indicated stimuli for 90 min. Pyrin S242 phosphorylation was assessed by Western blot analysis following immunoprecipitation. <u>Data information</u>: (A-E) Mean and SEM are shown. (B-C) each dot represents the value for one HD. (D-E) One experiment with technical triplicates representative of two (D) to three (E) independent experiments is shown.

Current knowledge places the activation of the pyrin inflammasome downstream of RhoA GTPases inhibition ^5, 8^. Yet, we observed no decrease in RhoA activity after etiocholanolone, a result that contrasted with the robust RhoA inhibition observed upon TcdA treatment (Fig. 4D, supplemental Fig. S3A). This observation suggests that steroid catabolites activate pyrin by a mechanism distinct from the one triggered by TcdA. We then investigated the pyrin domains required for steroid catabolite-mediated pyroptosis and IL-1β release. Pyrin proteins lacking either the pyrin (PYD) domain, the exon 2- encoded phosphorylated linker (PLD), the B-box (BBox), the coiled-coil (CC) or the B30.2 domains were stably expressed in U937 cells (Supplemental Fig. S3B). The resulting cell lines were treated with pregnanolone, etiocholanolone or TcdA. While the B30.2 domain was dispensable for TcdA-mediated response, steroid catabolites did not trigger release of IL-1β in absence of the B30.2 domain (Fig. 4E). Conversely, in absence of the phosphorylated linker domain, steroid catabolites triggered IL-1β release, further strengthening the evidence that the activation mechanism is primarily independent of step 1. The PYD, B-Box and coiled-coil domains were required for both steroid catabolites and TcdA responses. Furthermore, all the different cell lines responded similarly to NLRP3 stimulation by LPS + nigericin.

The difference in the B30.2 dependence was further validated by comparing pyrin serine 242 dephosphorylation after treatment with TcdA or steroid catabolites. Indeed, TcdA and UCN-01 triggered the dephosphorylation of both WT and ΔB30.2 pyrin proteins while etiocholanolone and pregnanolone only triggered pyrin dephosphorylation in the presence of the B30.2 domain (Fig. 4F). These experiments indicated that pyrin dephosphorylation happened in a B30.2-dependent manner specifically after addition of steroid catabolites.

Although the exact molecular mechanisms remain to be deciphered, these experiments revealed an activation mechanism that strongly differs from the one triggered by RhoA- inhibiting toxins. Indeed, pyrin activation by steroid catabolites is initiated in a B30.2- dependent manner, takes place in the absence of RhoA inhibition and does not require the phosphorylated linker domain, which includes the two serine residues required to launch TcdA/B-mediated pyrin responses ^10^.

### The response to steroid catabolites is specific to human pyrin

Mouse pyrin does not contain a B30.2 domain ^18^ but is a functional protein triggering inflammasome activation in response to RhoA-inhibiting toxins ^5^. In addition, the B30.2 domain is highly polymorphic in primates^19^. Since we observed a total dependence on the B30.2 domain for the response to steroid catabolites, we assessed whether pyrin proteins from other species could promote responsiveness to steroid catabolites. Etiocholanolone and pregnanolone did not trigger pyroptosis in U937 cells expressing either mouse or macaque pyrin (Fig. 5A, B and supplemental Fig. S4) while these cells underwent cell death in response to TcdA (Fig. 5C). These results were further confirmed by quantifying IL-1β secretion (Fig. 5D). The unresponsiveness of the murine pyrin inflammasome to etiocholanolone and pregnanolone was further validated in primary murine macrophages (Fig. 5E).

**Figure 5:**
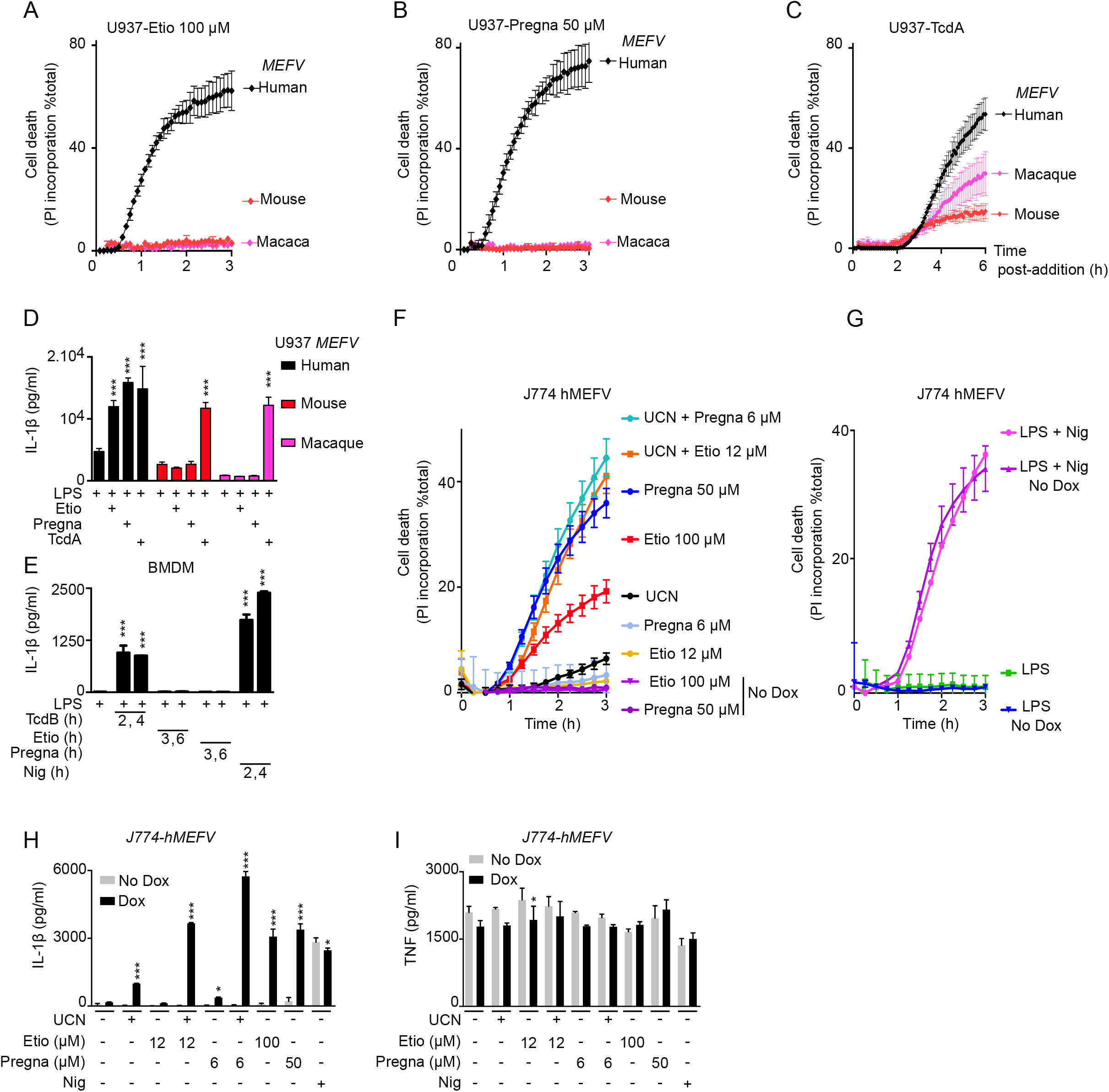
The human specificity of the response to steroid catabolites is intrinsic to the pyrin protein. (A-C) Doxycycline-induced U937 monocytes or (D) PMA-differentiated U937 macrophages expressing human (black), mouse (red) or *macaca fascicularis* (magenta) *MEFV* were treated with the indicated stimuli. (A-C) Propidium iodide (PI) incorporation was monitored every 5 min for 3 to 6 h. (D) IL-1β concentration in the supernatant was quantified at 3 h (Etio, Pregna) or 6 h (TcdA) post-addition. (E) WT bone-marrow derived macrophages (BMDM) were primed for 16 h with LPS (100ng/ml) and treated with TcdB (10ng/ml), etiocholanolone (100 μM), pregnanolone (50 μM), or nigericin (10 μg/mL). IL-1β concentration in the supernatant was quantified at the indicated time point post-compound addition. (F-I) J774 macrophages expressing or not (No Dox) human MEFV were treated with the indicated stimuli. (F-G) Propidium iodide (PI) incorporation was monitored every 5 min for 3 h. (H-I) Cells were primed for 3h with LPS before stimuli addition. (H) IL-1β and (I) TNF concentrations in the supernatant were quantified at 3 h post-addition. <u>Data information:</u> (A-I) one experiment representative of three independent experiments with mean and SEM of biological triplicates is shown. (D-E, H-I) One-way ANOVA with Sidak’s test was used, *: p<0.05; ***: p<0.001.

To test whether the lack of response of murine macrophages to steroid catabolites was intrinsic to the pyrin protein or linked to a more general defect, we expressed human pyrin in the murine macrophage cell line, J774. The expression of human pyrin was sufficient to recapitulate the responses seen in human cells (Fig. 5F, 5H). The response to NLRP3 stimulation was independent of human pyrin expression (Fig. 5G, H) while, as expected, TNF levels were not impacted by human pyrin expression (Fig. 5I). Altogether, these results demonstrate that the response to steroid catabolites is human-specific and that this species specificity is conferred, in an intrinsic manner, by the pyrin protein.

### Monocytes from FMF patients display a moderate increase in the response to steroid catabolites compared to HD

We then investigated whether FMF-associated mutations in *MEFV* exon 10 (altering the B30.2 domain) had an impact on the response of the pyrin inflammasome to steroid catabolites. U937 cells expressing three clearly pathogenic *MEFV* variants ^20^ were exposed to etiocholanolone (Fig. 6A and supplemental Fig. S5) and pregnanolone and EC50 were determined. While p.M680I and p.M694V mutations decreased these EC50 (reaching statiscal significance for p.M680I in the case of pregnanolone), the less severe p.V726A mutation^21^, significantly increased the EC50 of both etiocholanolone and pregnanolone. We then evaluated the steroid catabolites response of monocytes from FMF patients presenting at least one p.M680I or p.M694V mutation. A trend towards a faster and stronger cell death response was observed in response to steroid catabolites in monocytes from FMF patients compared to HD (Fig. 6B, C). No difference was observed in response to NLRP3 inflammasome engagement (Fig. 6B, C) while, as previously described^11, 17^, UCN-01-mediated pyroptosis was specifically observed in monocytes from FMF patients (Fig. 6B, C).

**Figure 6:**
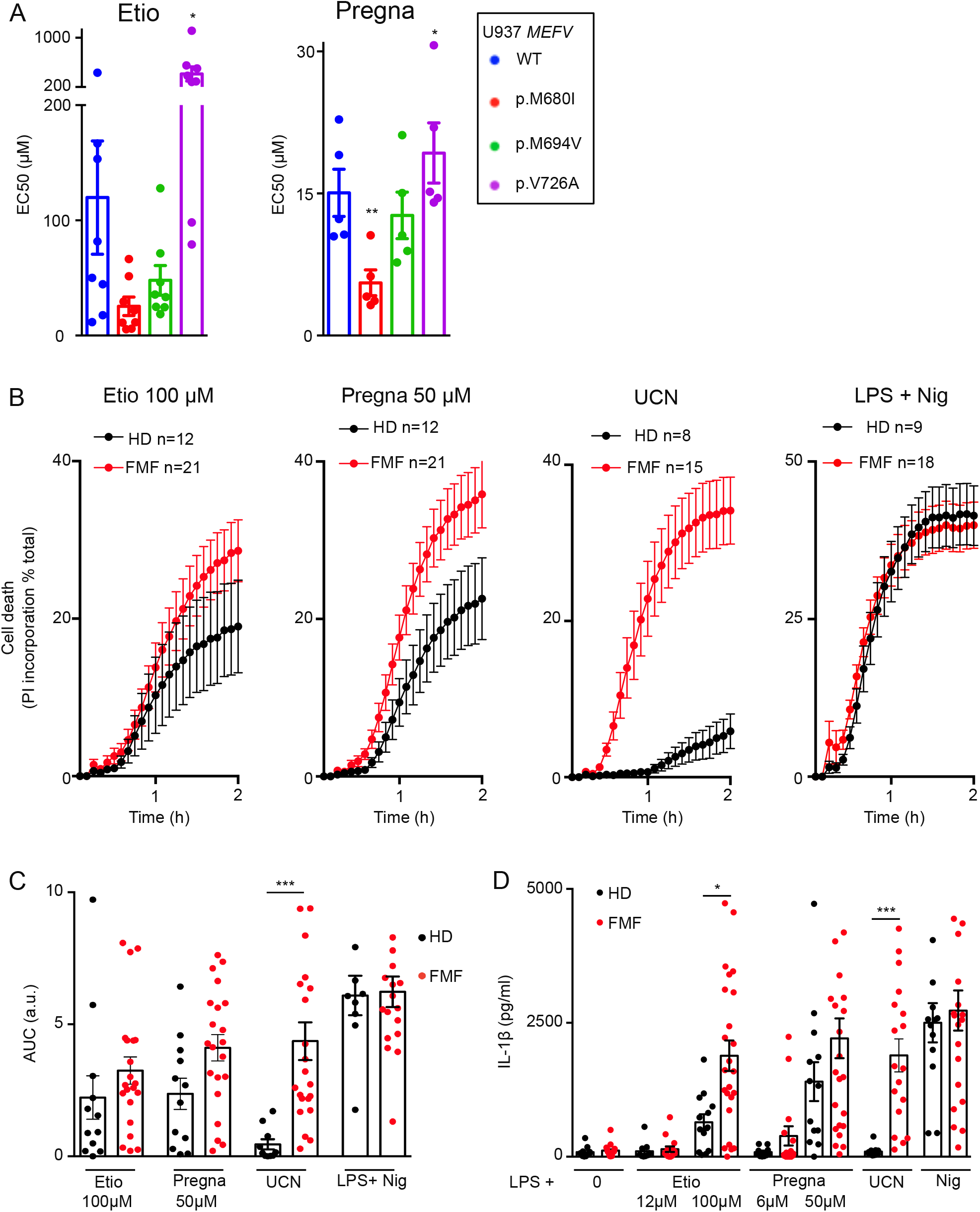
Monocytes from FMF patients display a moderately increased response to steroid catabolites compared to HD. (A) U937 expressing the indicated *MEFV* variant were exposed to the indicated steroid catabolite and the EC50 was determined at 3 h post-addition. (B) monocytes from HD (n=8-12) or FMF patients (n=15-21) were treated with the indicated stimuli. Propidium iodide (PI) incorporation was monitored every 5 min for 2 h. (C) The corresponding area under the curve (AUC) are shown. (D) IL-1β concentrations were determined at 3 h post addition of the indicated molecules. <u>Data information:</u> (A) Mean and SEM of five to eight independent experiments are shown. Each dot represents the mean value of a triplicate from one experiment. RM one-way ANOVA with Dunnet’s multiple comparisons test was performed. Etio *: p=0.018; Pregna *: p=0.033 **: p=0.0042. (B) Mean and SEM of 8 to 21 individuals are show as indicated, each one corresponding to the average of a biological triplicate. (C) Each dot corresponds to the mean AUC of one individual performed in triplicates, the bar represents the mean of 8 to 21 individuals. (D) Each dot corresponds to the mean IL-1β concentrations of one individual performed in triplicates, the bar represents the mean of 8 to 21 individuals. (C-D) One-way Anova with Sidak’s multiple comparison test was applied; *p=0.015; ***p<0.001.

FMF monocytes released on average 4.5-fold and 1.6-fold more IL-1β than HD monocytes in response to low or high concentrations of pregnanolone, respectively (Fig. 6D). In response to 100μM etiocholanolone, monocytes from FMF patients released significantly more IL-1β than HD’s monocytes (p=0.015). The difference in IL-1β concentrations were much stronger in response to UCN-01 (19.9-fold increase, p<0.001) while no difference was observed upon NLRP3 stimulation. Altogether, these results suggest that FMF patients display a moderate increase in steroid catabolite-induced inflammasome responses that could contribute to inflammatory flares and be dependent on the *MEFV* genotype.

### Monocytes from PAAND patients respond to low concentrations of steroid catabolites in the absence of step 1 activator

To assess the impact of PAAND-associated *MEFV* mutations on the response to steroid catabolites, we generated U937 cells expressing either of three reported PAAND mutations (p.S208C, p.S242R, p.E244K-supplemental Fig. S6A). Contrary to WT or p.M694V-expressing cell lines, all PAAND cell lines died quickly in response to low concentrations of pregnanolone or etiocholanolone (Fig. 7A, B). UCN-01 synergized with the low concentrations of steroid catabolites in p.S208C-expressing cells whereas it had no additional effect in p.S242R or p.E244K-expressing cells. Phosphorylation of serine residue 242 and interaction of 14-3-3 chaperone with the neighbouring residues may thus be more important to maintain pyrin inactive than phosphorylation of serine residue 208. This hypothesis is consistent with the fact that p.S242R and p.E244K mutations promote disease in a dominant manner while p.S208C does so in a recessive manner ^9,14,15^.

**Figure 7:**
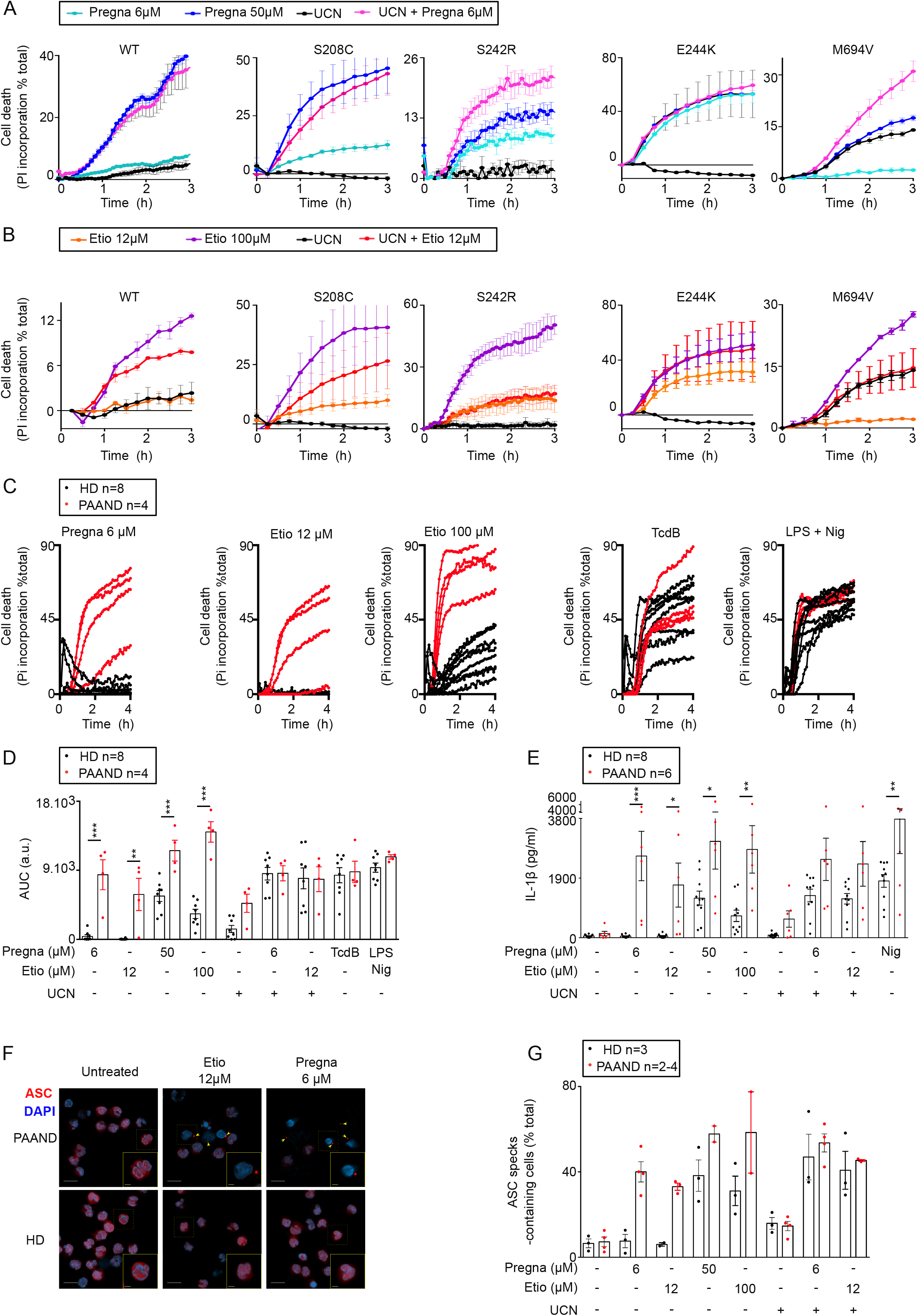
PAAND mutations confer hyper-responsiveness to steroid catabolites. (A-B) U937 cell lines expressing WT or the indicated PAAND MEFV variant were treated with the indicated stimuli. Propidium iodide (PI) incorporation was monitored every 15 min for 3 h. (C-G) monocytes from PAAND patient(s) (red, n=4-6) or HD (black, n=8) were treated with the indicated stimuli. (C) Propidium iodide (PI) incorporation was monitored every 15 min for 4 h. (D) The corresponding area under the curve (AUC) are shown. (E) IL-1β concentrations were determined in the cell supernatant after 3 h of LPS treatment followed by 1 h 30 of the indicated treatment. (F-G) ASC speck formation (indicated by yellow arrows) were monitored by immunofluorescence at 90 min post treatment. (F) Representative images are shown. The scale bars correspond to 10 and 2.5 μM, in the main pictures and insets, respectively. (G) Quantification is shown. <u>Data information:</u> (A-B) One experiment representative of three independent experiments is shown. (A-B) Mean and SEM or (C) mean of triplicates are shown. (D, E) Each dot represents the mean of a triplicate for one individual. The bar represents the mean +/- SEM. One way ANOVA with Sidak’s correction for multiple test was performed. (D) **: p=0.0011; ***: p=<0.001; a.u.: arbitrary units. (E) Etio 12 μM *:p=0.043; Pregna 50 μM *:p=0.016; Etio 100 μM **:p=0.0026; LPS Nig **:p=0.0064; ***: p=0.0001. (G) Each dot corresponds to the percentage of ASC specks based on more than 100 cells counted per condition. Each dot represents the value for one individual. The bar represents the mean +/- SEM.

We then assessed pyroptosis and IL-1β release in primary monocytes from PAAND patients from two independent families with heterozygous p.S242R mutation. At low concentration, pregnanolone and etiocholanolone triggered cell death (Fig. 7C, D), IL-1β release (Fig. 7D, E) and ASC specks formation (Fig. 7F, G). In contrast, none of these responses were observed in monocytes from HD. No differences were observed upon stimulation with TcdB or UCN-01 (in the presence or absence of low concentrations of steroid catabolites) (Fig. 7C-E, G). High concentrations of pregnanolone or etiocholanolone also triggered higher IL-1β release in PAAND patients’ monocytes compared to HD monocytes (Fig. 7E) possibly due to a faster and stronger inflammasome response (Fig. 7 D, supplemental Fig. S6B).

These results thus demonstrate that PAAND patients’ monocytes strongly respond to low doses of steroid catabolites suggesting that these molecules could contribute to inflammation in these patients and also to the distinct clinical features observed in PAAND and FMF patients.

## Discussion

The identification of steroid catabolites as step 2 triggers provides new insights into pyrin activation mechanisms. First, it confirms that the two steps can be activated independently. Indeed, low doses of sex steroid catabolites do not impact pyrin phosphorylation but the same doses trigger inflammasome activation in the presence of pyrin variants impaired for phosphorylation. Furthermore, the step 2 is dependent on the B30.2 domain, and independent of the phosphorylated linker domain, while the inverse applies to TcdA-mediated pyrin activation. Finally, a coupling mechanism likely exists between two steps since high doses of steroid catabolites trigger the full activation of pyrin. Of note, in this particular setting, what we initially termed “second step” is likely to be upstream of pyrin dephosphorylation.

In contrast to the TcdA/B responses, the response to steroid catabolites is human-specific. In mice, the lack of B30.2 domain partly explains the absence of response. Yet, we did not observe any response in bone-marrow derived macrophages from knock-in mice presenting the murine pyrin protein fused to the human B30.2 domain^22, 23^ (Supplemental Fig. S7). The resulting chimeric pyrin does not include the human coiled-coil domain likely explaining the lack of response of these cells. Indeed, the coiled-coil domain is required for steroid catabolite-mediated pyrin activation and is highly divergent between murine and human pyrin proteins (Supplemental Fig. S4). Surprisingly, the expression of *Macaca fascicularis* pyrin did not confer responsiveness to steroid catabolites either, despite the presence of similar coiled-coil and B30.2 domains. *MEFV* has been subjected to a strong evolutionary pressure in primates ^19^, possibly explaining the difference in steroid catabolites responsiveness between *M. fascicularis* and *Homo sapiens*.

Sex hormones are known to regulate immune responses and their variations during menstrual cycle and pregnancy correlate with profound modifications in local and systemic immune responses ^24–26^. Particularly, IL-1β and IL-18 levels fluctuate during menstrual cycle and pregnancy ^27–29^. Furthermore, flares in women with FMF are frequently associated with menstruation ^30–32^, which correspond to the peak of progesterone catabolism. While this correlation is appealing, demonstrating the pathophysiological role of pregnanolone (and/or etiocholanolone) remains challenging, especially due to the current lack of animal models and the complexity of the metabolome changes during menstrual cycles and pregnancy ^25^.

Progesterone and testosterone were completely inactive with regards to pyrin inflammasome activation. Similarly, all the modifications tested on the etiocholanolone or the pregnanolone backbones decreased inflammasome responses. We thus believe that human pyrin has specifically evolved in humans to sense these catabolites. Interestingly, Alimov and colleagues identified a synthetic molecule, BAA473, that shares the same steroid backbone and the same stereochemistry as etiocholanolone and pregnanolone and activates pyrin. In contrast to the molecules identified here, which are endogenous molecules, the relevance of BAA473 remains to be established although, theoretically, BAA473 could be generated from the secondary bile acid, deoxycholic acid^33^. BAA473-mediated pyrin inflammasome activation likely proceeds through an identical mechanism as steroid catabolites.

In addition to being sex hormones catabolites, pregnanolone and etiocholanolone are also neurosteroids that can be generated de novo in the central nervous system ^34^. Neurosteroids levels vary greatly, depending on the specific physiological situations, and can reach submicromolar to micromolar concentrations ^35^. Particularly, pregnanolone levels increase during psychological stress ^36, 37^, a condition known to promote flares in FMF patients ^32^. Whether etiocholanolone or pregnanolone could locally reach high enough concentrations to prime or activate the WT pyrin inflammasome in a particular neurological environment (and contribute to neuroinflammation) is unclear at the moment. Interestingly, we observed that at least one *MEFV* mutation (p.P373L) confers responsiveness to nanomolar concentrations of steroid catabolites (Fig. S8) indicating that physiological concentrations can modulate pyrin inflammasome activation. All other neurosteroids tested (pregnenolone and pregnenolone sulfate) were inactive, and we could not identify receptors upstream of the pyrin inflammasome to explain the pyroptotic effect of pregnanolone and or etiocholanolone. The pyrin B30.2 domain contains a hydrophobic pocket that has been hypothesized to bind a ligand. Similarly, the butyrophilin 3A1 B30.2 domain displays a pocket that accommodates microbial-derived phosphoantigens resulting in the activation of gamma delta T cells ^38^. It is thus tempting to speculate that pregnanolone and etiocholanolone could directly bind the pyrin B30.2 domain to activate the inflammasome.

Importantly, experiments performed in the late 50s have demonstrated the fast and potent pyrogenic activity of steroids of endogenous origin, including etiocholanolone and pregnanolone upon injection in human volunteers ^39, 40^ (Supplemental Fig. S9). These historical experiments thus validate these steroid catabolites as potent in vivo inflammation inducers and our results strongly suggest that activation of the pyrin inflammasome was at the origin of this enigmatic “Steroid fever” ^41^.

## Methods

### Ethics statement

The study was approved by the French Comité de Protection des Personnes (CPP,#L16-189) and by the French Comité Consultatif sur le Traitement de l’Information en matière de Recherche dans le domaine de la Santé (CCTIRS, #16.864) and the Leuven/Onderzoek Ethic committee (#S58600). The authors observed a strict accordance to the Helsinki Declaration guidelines. HD blood was provided by the Etablissement Français du Sang in the framework of the convention #14-1820, after receiving written informed consent from all donors.

### Subjects

All FMF patients fulfilled the Tel Hashomer criteria for FMF and had at least one mutation in the *MEFV* gene. PAAND patients all bear heterozygous p.S242R mutation. Three patients have been previously reported ^42^ while three patients (1 adult 2 children) were newly identified by Pr. Tran (CHU Nîmes). The potential carriage of *MEFV* mutations in HD was not assessed. Blood samples from HD were drawn on the same day as patients.

### Reagents

Etiocholanolone (3α-hydroxy-5β-androstan-17-one, #R278572), Testosterone (#86500), Androsterone (3α-hydroxy-5α -androstan-17-one, #31579), 3β-hydroxy-5β-androstan-17-one (R213691), Progesterone (#P8783), Pregnanolone (5- beta-pregnan-3α-ol, 20-one, #P8129), cortisol (#C-106), UCN-01 (#U6508), Doxycycline (#D9891) were from Sigma. Pregnanolone (5-beta-pregnan-3α-ol, 20-one, #P8150-000), 11-one: (5-beta-pregnan-3α-ol-11, 20-dione, #P7850-000), 11αOH: (5β-pregnan-3α, 11α-diol-20-one, #P6400-00), 11βOH: (5-β-pregnan-3-α, 11β-diol-20-one, #P6420-000), Hemisuccinate: (5-β-pregnan-3-α, 21-diol-20-one, 21 hemisuccinate, #P6944-000), 21OH: (5-β-pregnan-3-α, 21-diol-20-one, #P6920-000), 17OH (5-β-pregnan-3-α, 17 diol-20-one, #P6570-000), 20H (5-β-pregnan-3-α-ol, #P7800-000), Sulphate (5β- pregnan-3α-ol-20-one sulphate, sodium salt, #P8168-000) were from Steraloids. Tetrahydrocortisol (#T293370) was from Toronto Research Chemicals. LPS-EB Ultrapure (#tlrl-3pelps), Nigericin (#tlrl-nig) were from Invivogen. TcdB was from Abcam (#ab124001). TcdA was purified from *Clostridium difficile* VPI10463 strain, as previously described ^43, 44^.

### Chemical library screening

All robotic steps were performed on a Tecan Freedom EVO platform. Compounds from the Prestwick Chemical Library→ (https://www.prestwickchemical.com/screening-libraries/prestwick-chemical-library/) were evaluated at a 1:1,000 dilution of the original stock: 2 mg/mL corresponding to 6.32 ± 2.8 mM for plates 1 to 14 and 10 mM for the last plate. 1 µL of DMSO solutions was spiked into dry well of F-bottom clear cell culture treated 96-wells plates (Greiner Bio One), with columns 1 and 12 devoted to controls and used to calculate the Z’-factor. U937 cells expressing p.S242R *MEFV* were treated or not (counterscreen) with doxycycline (1 μg/mL) for 16 h, centrifuged and seeded at 10^5^ cells per well (100μL final volume) in RPMI 1640 without phenol red, 10% FCS, 1mM Hepes, 1% PSA, 1mM Glutamine, in the presence of propidium iodide at 5 μg/ml. Following 90 min of incubation at 37°C, fluorescence intensity (excitation wavelength at 535 nm and emission wavelength at 635 nm) corresponding to propidium iodide incorporation was measured on a microplate reader (Infinite M1000, Tecan). The average Z’ value was 0.67 ± 0.12, indicating a robust and reliable assay. Mean fluorescence + 3SD was retained as a threshold. Compounds triggering cell death only in the presence of doxycycline were considered as pyrin-specific and defined as hits.

### Monocyte and neutrophil isolation

Blood was drawn in heparin-coated tubes and kept at room temperature overnight. Monocytes were isolated as previously reported ^11^. Briefly, peripheral blood mononuclear cells (PBMCs) and neutrophils were isolated by density-gradient centrifugation. Monocytes were further isolated by magnetic positive selection using CD14 MicroBeads (Miltenyi Biotec) following manufacturer’s instructions. Neutrophils were separated from red blood cells (RBCs) using Dextran (Sigma, #31392), residual RBCs were lysed with ice cold bi-distilled water and contaminating CD14^+^ monocytes were excluded using CD14 MicroBeads. Live cells were enumerated by flow cytometry (BD Accuri C6 Flow Cytometer®).

### Bone-marrow derived macrophages

Bone-marrow derived macrophages from C57BL6/J (Charles River) or *MEFV*^M694VKI 23^ mice were obtained in the framework of the ethical approval ENS_2012_061 (CECCAPP, Lyon, France). Standard protocols were used as previously reported ^45^.

### Cell lines and Genetic manipulation

The human myeloid cell line U937 was grown in RPMI 1640 medium with glutaMAX-I supplemented with 10% (vol/vol) FCS, 2 mM L-glutamine, 100 IU/mL penicillin, 100 μg/mL streptomycin (ThermoFischer Scientific). *Casp1*^KO^ and *GSDMD*^KO^ cell lines, U937 cell lines expressing WT, p.S208C, p.S242R, p.M694V, p.M680I under the control of a doxycycline-inducible promoter, have been previously described^11, 46^. p.[V726A], p.[E244K], ΔPYD, ΔPLD, ΔB-Box, ΔCoiled-coil, ΔB30.2 *MEFV* were generated by mutagenesis of the pENTR1A-3xFlag *MEFV* using primers presented in supplemental table S1, pfu ultra II Fusion high fidelity polymerase (Agilent) followed by digestion of the parental plasmid using Dpn1 restriction enzyme. The resulting plasmids were validated by sequencing and the mutated *MEFV* constructs were transferred into the GFP-expressing plasmid pINDUCER21 ^47^ under the control of a doxycycline-inducible promoter using LR recombinase (Invitrogen). Lentiviral particles were produced in 293T cells using pMD2.G and psPAX2 (from Didier Trono, Addgene plasmids #12259 and #12260), and pINDUCER-21 plasmids. U937 cells were transduced by spinoculation and selected at day 4 post-transduction based on GFP expression on an Aria cell sorter and maintained polyclonal. Pyrin expression was induced by treatment with doxycycline (1 μg.mL^-1^) for 16 h before stimulation. All parental cell lines were tested for mycoplasma contamination.

### Inflammasome activation

For cytokine quantification, primary monocytes were seeded in 96-well plates at 5x10^3^ cells/well, in RPMI 1640, GlutaMAX medium (Thermofisher) supplemented with 10% foetal calf serum (Lonza) and incubated for 3 hours in the presence of LPS (10 ng/ml, Invivogen). Primary monocytes were then treated for 1 h 30 with nigericin (5 μM, Invivogen); UCN-01 (12.5 μM, Sigma), TcdB (125 ng/ml, Abcam) or steroid catabolites at the indicated concentrations. When indicated, monocytes were treated with colchicine (1 μM, Sigma), nocodazole (5 μM, Sigma), VX-765 (25 μM, Invivogen) or MCC950 (10μM, Adipogen AG-CR1-3615) 30 minutes before addition of steroid catabolites, UCN-01, TcdB or Nigericin. Following the incubation, cells were centrifuged, and supernatants were collected.

To assess cytokine release, 8 x 10^4^ U937 cells per well of a 96 wells plate were exposed to 100 ng.mL^-1^ of phorbol 12-myristate 13-acetate (PMA; InvivoGen) for 48 h and primed with LPS at 1 μg.mL^-1^ for 3 h. When applicable, nigericin was used at 50 μg.mL^-1^. Supernatant was collected at 3 h post treatment. Levels of IL-1β or TNF in cell supernatants were quantified by ELISA (R&D Systems).

### ASC specks Immunofluorescence

Monocytes were fixed with paraformaldehyde 2% for 20 min before spreading onto poly-lysine adhesion slides (Thermo Scientific^TM^) using the Cytospin3 (Shandon) 5 min at 450 rpm. Following permeabilisation with Triton X100 (0.1% in PBS), cells were stained using anti-ASC (Santa Cruz, sc22514R, 4 μg.mL^-1^), Alexa594-goat anti rabbit antibodies (Invitrogen, A-110088, 10 μg.mL^-1^) and DAPI (100 ng.mL^-1^). ASC specks were visualized on the Zeiss LSM800 confocal microscope. Quantification was performed on 10 fields per sample.

### Real time cell death and EC50 calculation

For real time cell death assays, monocytes and U937 cells were seeded at 2 or 5 x 10^4^ per well of a black 96 well plate (Costar, Corning), respectively, in the presence of propidium iodide (PI, Sigma) at 5 μg/ml. Three technical replicates per conditions were done. Real time PI incorporation was measured every 5 to 15 min immediately post-stimuli addition on a fluorimeter (Tecan) using the following wavelengths: excitation 535 nm (bandwidth 15 nm); emission 635 nm (bandwidth 15 nm) ^48, 49^. Cell death was normalized using PI incorporation in cells treated with triton X100 for 15 min (=100% cell death) and PI incorporation at each time point in untreated cells (0% cell death). As a further correction, the first time point of the kinetics was set to 0. The areas under the curve were computed using the trapezoid rule (Prism 6; GraphPad). To calculate the EC50 (Half maximal effective concentration), the normalized cell death at 3 h post-compound addition was used. To compare different cell lines, butyrate (1mM) was added for 16 h (in the meantime as doxycycline) to revert transgene silencing ^50^. The different concentrations were log-transformed, and a non-linear regression was applied using the Log (agonist) vs. normalized response-variable slope model (Prism 6; GraphPad). The least squares (ordinary) fitting method was applied.

### RhoA activity

RhoA activity was determined by G-LISA (Cytoskeleton) following manufacturer’s instructions.

### Immunoprecipitation and Immunoblot analysis

Cells were lysed in 25mM Tris HCl, 150mM NaCl, 1mM EDTA and 0.1% NP-40 buffer containing Mini Protease Inhibitor Mixture (Roche) and sodium fluoride (Sigma) by a quick freezing and thawing step. Flag-Pyrin was immuno-precipitated using anti Flag M2 affinity gel (Sigma). Proteins were separated by SDS/PAGE on precast 4-15% acrylamide gels (Bio-rad) and transferred to TransBlot® Turbo™ Midi-size PVDF membranes (Bio-rad). Antibodies used were mouse monoclonal anti-FLAG® (Sigma-Aldrich, clone M2; 1:1,000 dilution), anti-Pyrin (Adipogen, AL196, 1: 1,000 dilution), anti-phospho S242 Pyrin (Abcam, ab200420; 1:1,000 dilution)^10^, anti-human Caspase-1 (Santa Cruz, sc515, 1: 1,000 dilution), anti-human GSGMD (sigma, HPA044487, 1: 1,000 dilution), anti-human IL-1β (Cell signalling, #12703, 1: 1,000 dilution). Cell lysates were reprobed with a mouse monoclonal antibody anti-β-actin (clone C4, Millipore; 1:5,000 dilution).

### Statistical analysis

Normality was verified using D’Agostino & Person omnibus normality test or Kolmogorov-Smirnov test with Dallal-Wilkinson-Lille for P value if the number of values was too small for the former test. Gaussian distribution was assumed for technical triplicates. Unmatched normalized values were analysed by Ordinary one-way ANOVA with Sidak’s multiple comparisons test. When normality could not be verified, matched values were analysed by the Friedman test, with Dunn’s correction or using Sidak’s multiple comparisons test. Normal matched values were analysed with RM one-way ANOVA, with the Greenhouse-Geisser correction and Dunnett correction for multiple comparisons. Unmatched values, for which normality could not be verified, were analysed using Kruskal-Wallis analysis with Dunn’s correction.

## Acknowledgments

We warmly thank the patients and their family for their involvement in this project. This work was performed in the framework of the Centre National de Reférence RAISE. We acknowledge the contribution of SFR Biosciences (UMS3444/CNRS, US8/Inserm, ENS de Lyon, UCBL) Platim, cytometry and PBES facilities. We acknowledge the contributions of the CELPHEDIA Infrastructure (http://www.celphedia.eu/), especially the centre AniRA in Lyon. We acknowledge the contribution of the Etablissement Français du Sang Auvergne - Rhône-Alpes, Mathieu Gerfaud-Valentin, Emmanuelle Weber, Agnès Duquesne, Marine Fouillet-Desjonquères (Lyon university hospital), Marion Delplanque (Tenon Hospital) for patient recruitment. We thank Pr Etienne Merlin (CHU Clermont-Ferrand), Emmanuel Lemichez (Pasteur institute, Paris), TH team and ImmunAID members for advice and stimulating discussions. **Funding:** This work is supported by an ANR grant (FMFgeneToDiag). This project has received funding from the European Union’s Horizon 2020 research and innovation programme under grant agreement No 779295.

## Data availability

All data are available in the current manuscript or the supplemental information.

---

Figure S1: Structure of the steroid molecules tested in this study

Figure S2: The response to steroid catabolites is conserved in neutrophils and is caspase-1-dependent, and NLRP3-independent in monocytes.

(A) Propidium iodide incorporation in neutrophils from one HD treated with pregnanolone or etiocholanolone was monitored every 5 min for 2 h. (B-C) HD monocytes (n=3) were treated with LPS for 2 h 30 followed by addition or not of (B) the caspase-1 inhibitor VX-765, (C) the NLRP3 inhibitor (MCC950) and 30 min later of the indicated molecules. IL-1β concentrations were determined in the cell supernatant 1 h 30 after the final treatment.

Data information: (A) One experiment representative of two independent experiments is shown. Mean and SEM of a technical triplicates are shown. Cell death was normalized using untreated neutrophils (0%) and Triton X100-treated neutrophils (100%). (B-C): Each dot represents the value of one HD (mean of a triplicate). The bar represents the mean +/- SEM of 3 HD values. Matched one-way ANOVA with Sidak’s multiple comparisons test was performed to compare untreated vs. VX-765-treated samples. *: p<0.05.

Supplemental Fig. S3: Kinetics of TcdA and etiocholanolone-mediated cell death (related to Fig. 4D) and Western blot analysis of pyrin deletion mutants.

(A) U937 cells expressing WT pyrin (in the presence of doxycycline) were treated with etiocholanolone or TcdA. Cell death was monitored by following propidium iodide incorporation every 5 min for 7 h. The dotted vertical lines indicate the time points at which U937 cells were collected in a parallel experiment to assess RhoA inhibition (see Fig. 3D; 1, 2, 3 h for etiocholanolone and 3, 6 h for TcdA treatment).(B) U937 cell lines expressing the indicated 3xFlag-*MEFV* variants were analyzed by Western blot in the presence or absence of doxycycline (Dox). Actin was used as a loading control.

Supplemental Fig. S4: Alignment of human, mouse and macaca fascicularis pyrin proteins and validation of cell lines with ectopic expression of human, mouse or macaque pyrin.

(A) The PYD, the two critical serine residues, the B-Box, Central Helical scaffold (CHS) and B-30.2 domains are shown. (B) Western blot analysis of U937 (left panel) or J774 (right panel) cells expressing 3xFlag-pyrin from the indicated species. Cells were treated or not with doxycycline (DOX) and cell lysates were analysed with anti-Flag (top panel) or anti-actin (bottom panel) antibodies.

Supplemental Fig. S5: Western blot analysis of U937 cell expressing p.V726A pyrin.

U937 cell lines expressing the indicated pyrin variants were treated or not with doxycycline (Dox) and cell lysates were analysed with anti-pyrin (top panel) or anti-actin (bottom panel) antibodies. Cell lines expressing WT, M694V and M680I pyrin variants have been previously characterized ^11^.

Supplemental Fig. S6: PAAND patients monocytes present a strong increase in steroid catabolites responses compared to HD.

(A) Western blot analysis of U937 cell expressing p.E244K pyrin variant. U937 cell lines expressing the indicated pyrin protein were treated or not with doxycycline (Dox) and cell lysates were analysed with anti-Pyrin (top panel) or anti-actin (bottom panel) antibodies. Cell lines expressing WT, S208C and S242R pyrin variants have been previously characterized ^11^. (B) Monocytes from PAAND patients (red, n=4) or HD (black, n=8) were treated with pregnanolone (50 μM). Cell death/ propidium iodide incorporation was monitored in real time every 15 min for 4 h. Each dot corresponds to the average of a triplicate for one individual.

Supplemental Fig. S7: Bone marrow derived macrophages (BMDM) from human B30.2 knock-in mice do not respond to steroid catabolites.

BMDM from *mefv*^KI hB^^30.2p.M694V^ mice harbouring human B30.2 domain in fusion with murine pyrin protein were treated with the indicated steroid molecules for 1 h followed by addition (or not) of staurosporine (1 μM) for 3 h. IL-1β concentrations were determined in the cell supernatant 4 h after steroid addition. Androsterone (Andro) was used as a negative control based on Fig. 2C results. Etiocholanolone and pregnanolone treatment did not differ from androsterone treatment in the presence or absence of the PKC superfamily inhibitor, staurosporine, used here to trigger pyrin step 1. One way ANOVA with Sidak’s multiple comparisons test was performed. N.S.: not significant.

Supplemental Fig. S8: p.P373L *MEFV* variant confers responsiveness to nanomolar concentrations of steroid catabolites.

U937 cells expressing the indicated *MEFV* variant (green p.P373L; blue WT) were treated with doxycycline for 16 h followed by addition of pregnanolone or etiocholanolone at various concentrations. Cell death was measured at 3 h post-addition.

Data information: One experiment representative of three independent experiment is shown. Mean and SEM of triplicates are shown. Non-linear regression curve computed using least squares fit method is shown.

Supplemental Fig. S9: Etiocholanolone and pregnanolone trigger steroid fever in humans.

Figure from ^40^. Historical evidence that intravenous injection of etiocholanolone (top panel) or pregnanolone (bottom panel) trigger fever in humans. Results from 4 individuals healthy volunteers are shown.

Table S1: Oligonucleotides used in this study

