## Supplemental Fig for "Steroid hormone catabolites activate the pyrin inflammasome through a non-canonical mechanism"

| Name | Structure | Name | Structure |
|---|---|---|---|
| Etiocholanolone<br>(3α-hydroxy 5β-androstan-17one) | 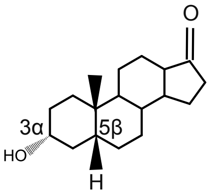 | 5β-Pregnan-3α,11α-diol -20-one | 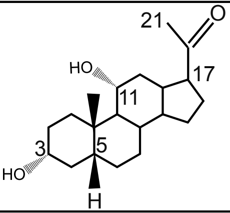 |
| Pregnanolone<br>(3α-hydroxy 5β-Pregnan-20one) | 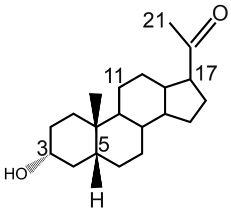 | 5β-Pregnan-3α,11β-diol -20-one | 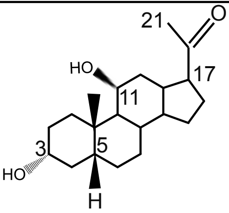 |
| Testosterone | 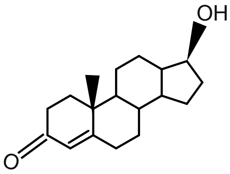 | 3α-hydroxy 5β-Pregnan-11,20-dione | 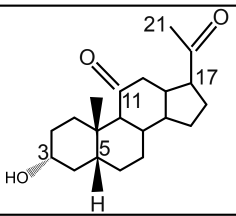 |
| Progesterone | 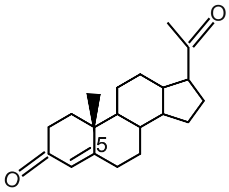 | 5β-Pregnan-3α, 17diol-20-one | 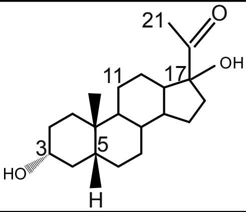 |
| Cortisol | 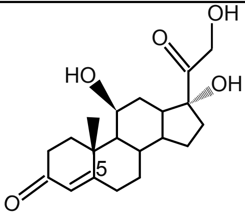 | 5β-Pregnan-3α-ol | 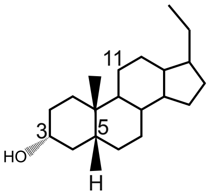 |
| Tetrahydro-cortisol | 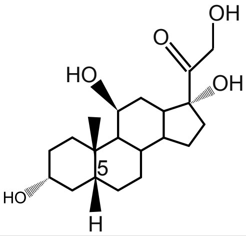 | 5β-Pregnan-3α,21-diol-20-one | 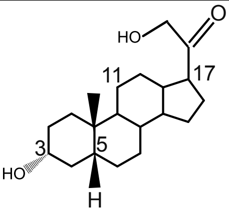 |
| Pregnanolone-sulfate | 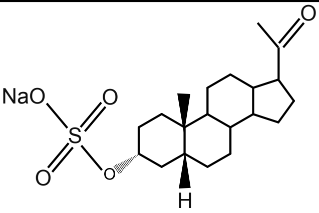 | 5β-Pregnan-3α,21-diol-20-one<br>21-hemisuccinate | 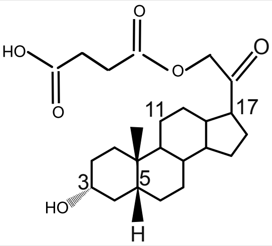 |
| 3β-hydroxy 5β-androstan-17one | 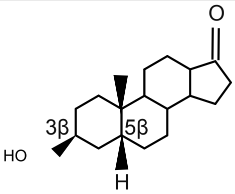 | Androsterone | 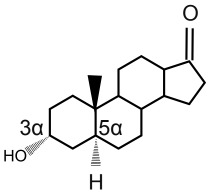 |

Fig. S1

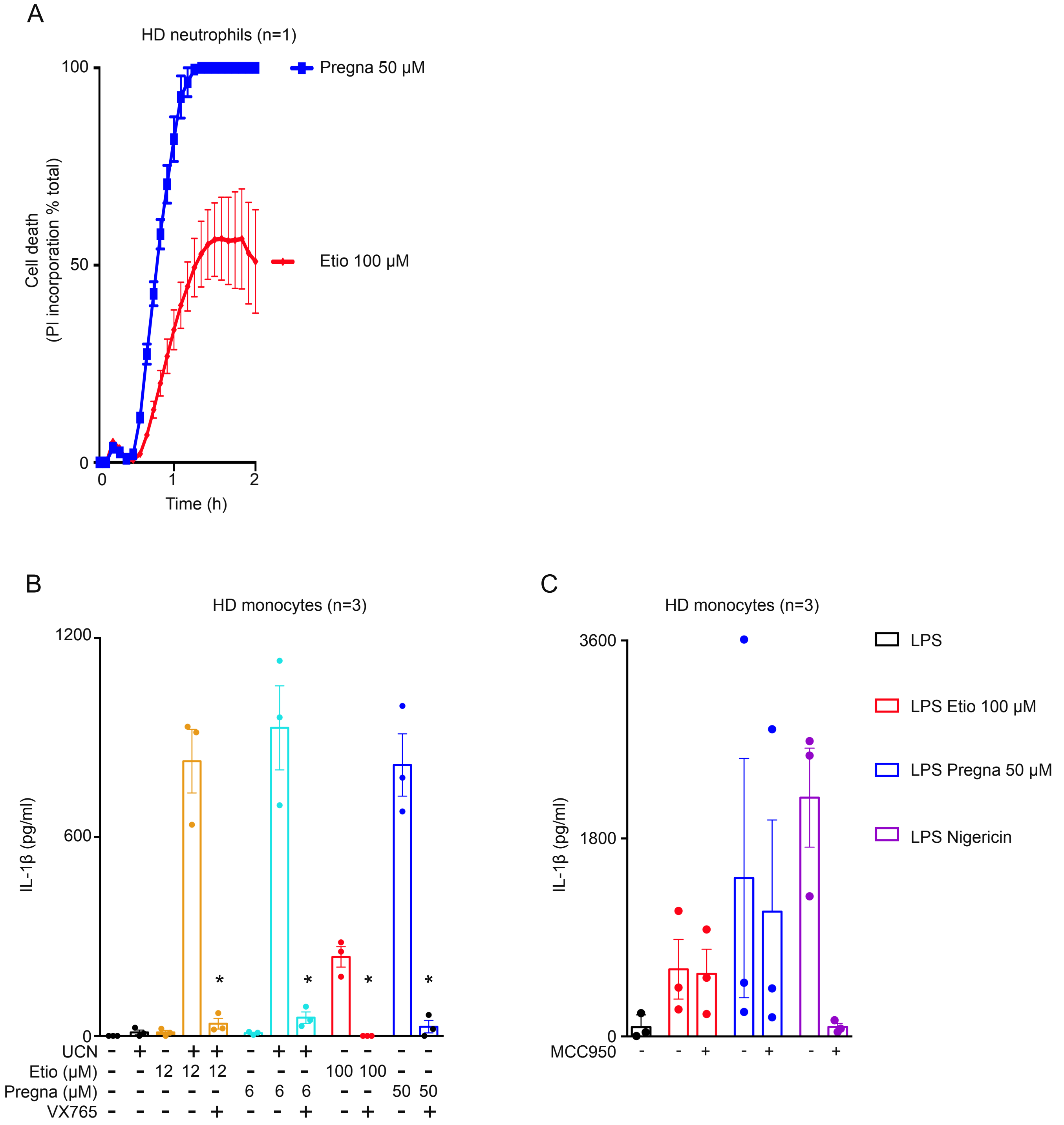
A
HD neutrophils (n=1)
Cell death (PI incorporation % total)
Time (h)
Pregna 50 μM
Etio 100 μM
B
HD monocytes (n=3)
IL-1β (pg/ml)
UCN
Etio (μM)
Pregna (μM)
VX765
C
HD monocytes (n=3)
IL-1β (pg/ml)
MCC950
LPS
LPS Etio 100 μM
LPS Pregna 50 μM
LPS Nigericin

Fig. S2

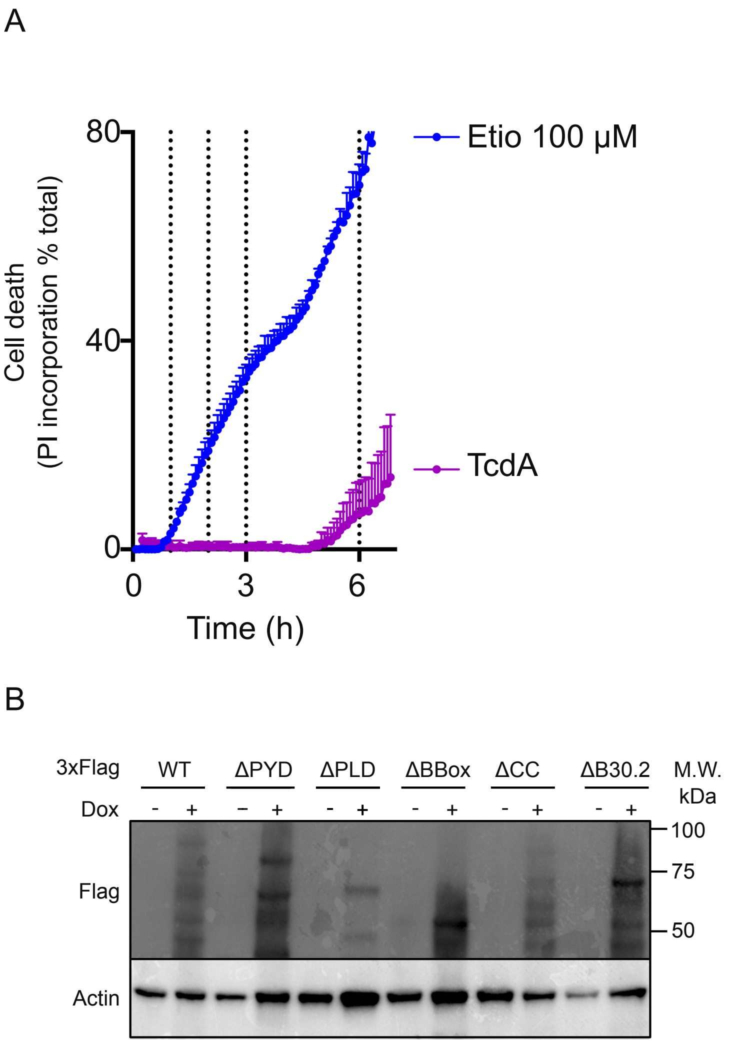
A
80
40
0
0
3
6
Cell death
(PI incorporation % total)
Time (h)
Etio 100 μM
TcdA
B
3xFlag
WT
ΔPYD
ΔPLD
ΔBBox
ΔCC
ΔB30.2
M.W.
kDa
Dox
- + - + - + - + - + - +
Flag
Actin
100
75
50

Fig S3

A

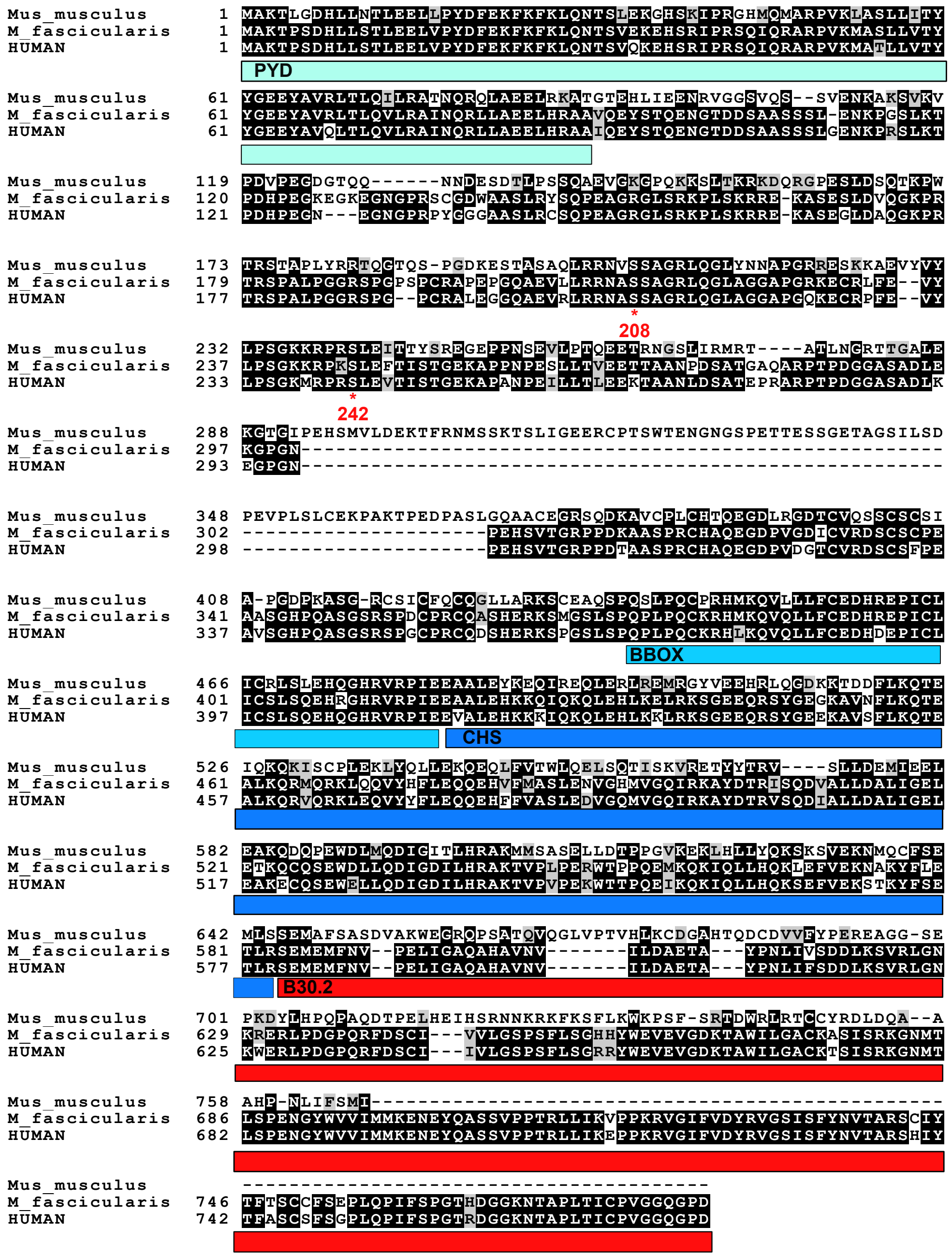
Mus_musculus 1 MAKTLGDHLLNTLEELLPYDFEKFKFKLQNTSLEKGHSKIPRGHMQMARPVKLASLLITY
M_fascicularis 1 MAKTPSDHLLSTLEELVPYDFEKFKFKLQNTSVEKEHSRIPRSQIQRARPVKMASLLVTY
HUMAN 1 MAKTPSDHLLSTLEELVPYDFEKFKFKLQNTSVQKEHSRIPRSQIQRARPVKMATLLVTY
PYD
Mus_musculus 61 YGEEYAVRLTLQILRATNQRQLAEELRKATGTEHLIEENRVGGSVQS--SVENKAKSVKV
M_fascicularis 61 YGEEYAVRLTLQVLRAINQRLLAEELHRAAVQEYSTQENGTDDSAASSSL-ENKPGSLKT
HUMAN 61 YGEEYAVQLTLQVLRAINQRLLAEELHRAAIQEYSTQENGTDDSAASSSLGENKPRSLKT
Mus_musculus 119 PDVPEGDGTQQ------NNDESDTLPSSQAEVGKGPQKKSLTKRKDQRGPESLDSQTKPW
M_fascicularis 120 PDHPEGKEGKEGNGPRSCGDWAASLRYSQPEAGRGLSRKPLSKRRE-KASESLDVQGKPR
HUMAN 121 PDHPEGN---EGNGPRPYGGGAASLRCSQPEAGRGLSRKPLSKRRE-KASEGLDAQGKPR
Mus_musculus 173 TRSTAPLYRRTQGTQS-PGDKESTASAQLRRNVSSAGRLQGLYNNAPGRRESKKAEVYVY
M_fascicularis 179 TRSPALPGGRSPGPSPCRAPEPGQAEVLLRRNASSAGRLQGLAGGAPGRKECRLFE--VY
HUMAN 177 TRSPALPGGRSPG--PCRALEGGQAEVRLRRNASSAGRLQGLAGGAPGQKECRPFE--VY
*
208
Mus_musculus 232 LPSGKKRPRSLEITTYSRFGEPPNSEVLPTQEETRNGSLIRMRT----ATLNGRTTGALE
M_fascicularis 237 LPSGKKRPKSLEFTISTGEKAPPNPESLLTVEETTAANPDSATGAQARPTPDGGASADLE
HUMAN 233 LPSGKMRPRSLEVTISTGEKAPANPEILLTLEEKTAANLDSATEPRARPTPDGGASADLK
*
242
Mus_musculus 288 KGTGIPEHSMVLDEKTFRNMSSKTSLIGEERCPTSWTENGNGSPETTESSGETAGSILSD
M_fascicularis 297 KGPGN-------------------------------------------------------
HUMAN 293 EGPGN-------------------------------------------------------
Mus_musculus 348 PEVPLSLCEKPAKTPEDPASLGQAACEGRSQDKAVCPLCHTQEGDLRGDTCVQSSCSCSI
M_fascicularis 302 ---------------------PEHSVTGRPPDKAASPRCHAQEGDPVGDICVRDSCSCPE
HUMAN 298 ---------------------PEHSVTGRPPDTAASPRCHAQEGDPVDGTCVRDSCSFPE
Mus_musculus 408 A-PGDPKASG-RCSICFQCQGLLARKSCEAQSPQSLPQCPRHMKQVLLLFCEDHREPICL
M_fascicularis 341 AASGHPQASGSRSPDCPRCQASHERKSMGSLSPQPLPQCKRHMKQVQLLFCEDHREPICL
HUMAN 337 AVSGHPQASGSRSPGCPRCQDSHERKSPGSLSPQPLPQCKRHLKQVQLLFCEDHDEPICL
BBOX
Mus_musculus 466 ICRLSLEHQGHRVRPIEEAALEYKEQIREQLERLREMRGYVEEHRLQGDKKTDDFLKQTE
M_fascicularis 401 ICSLSQEHRGHRVRPIEEAALEHKKQIQKQLEHLKELRKSGEEQRSYGEGKAVNFLKQTE
HUMAN 397 ICSLSQEHQGHRVRPIEEVALEHKKKIQKQLEHLKKLRKSGEEQRSYGEEKAVSFLKQTE
CHS
Mus_musculus 526 IQKQKISCPLEKLYQLLEKQEQLFVTWLQELSQTISKVRETYYTRV----SLLDEMIEEL
M_fascicularis 461 ALKQRMQRKLQQVYHFLEQQEHVFMASLENVGHMVGQIRKAYDTRISQDVALLDALIGEL
HUMAN 457 ALKQRVQRKLEQVYYFLEQQEHFFVASLEDVGQMVGQIRKAYDTRVSQDIALLDALIGEL
Mus_musculus 582 EAKQDQPEWDLMQDIGITLHRAKMMSASELLDTPPGVKEKLHLLYQKSKSVEKNMQCFSE
M_fascicularis 521 ETKQCQSEWDLLQDIGDILHRAKTVPLPERWTPPQEMKQKIQLLHQKLEFVEKNAKYFLE
HUMAN 517 EAKECQSEWELLQDIGDILHRAKTVPVPEKWTTPQEIKQKIQLLHQKSEFVEKSTKYFSE
Mus_musculus 642 MLSSEMAFSASDVAKWEGRQPSATQVQGLVPTVHLKCDGAHTQDCDVVFYPEREAGG-SE
M_fascicularis 581 TLRSEMEMFNV--PELIGAQAHAVNV-------ILDAETA---YPNLIVSDDLKSVRLGN
HUMAN 577 TLRSEMEMFNV--PELIGAQAHAVNV-------ILDAETA---YPNLIFSDDLKSVRLGN
B30.2
Mus_musculus 701 PKDYLHPQPAQDTPELHEIHSRNNKRKFKSFLKWKPSF-SRTDWRLRTCCYRDLDQA--A
M_fascicularis 629 KRERLPDGPQRFDSCI---VVLGSPSFLSGHHYWEVEVGDKTAWILGACKASISRKGNMT
HUMAN 625 KWERLPDGPQRFDSCI---IVLGSPSFLSGRRYWEVEVGDKTAWILGACKTSISRKGNMT
Mus_musculus 758 AHP-NLIFSMI-------------------------------------------------
M_fascicularis 686 LSPENGYWVVIMMKENEYQASSVPPTRLLIKVPPKRVGIFVDYRVGSISFYNVTARSCIY
HUMAN 682 LSPENGYWVVIMMKENEYQASSVPPTRLLIKEPPKRVGIFVDYRVGSISFYNVTARSHIY
Mus_musculus ----------------------------------------
M_fascicularis 746 TFTSCCFSEPLQPIFSPGTHDGGKNTAPLTICPVGGQGPD
HUMAN 742 TFASCSFSGPLQPIFSPGTRDGGKNTAPLTICPVGGQGPD

B

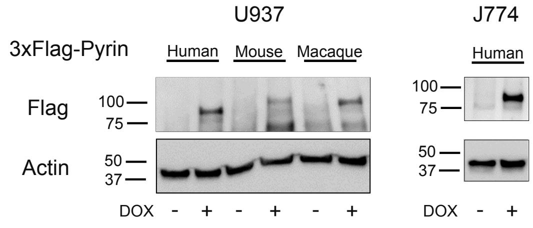
U937
J774
3xFlag-Pyrin
Human
Mouse
Macaque
Human
Flag
100
75
100
75
Actin
50
37
50
37
DOX - + - + - +
DOX - +

Fig. S4

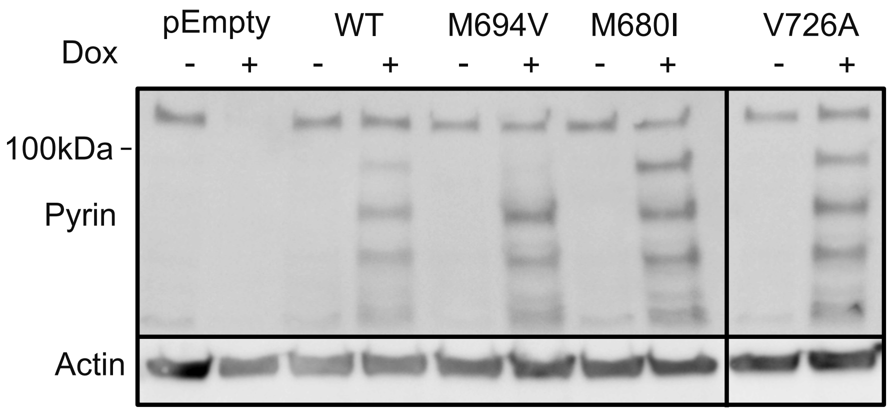


Fig. S5

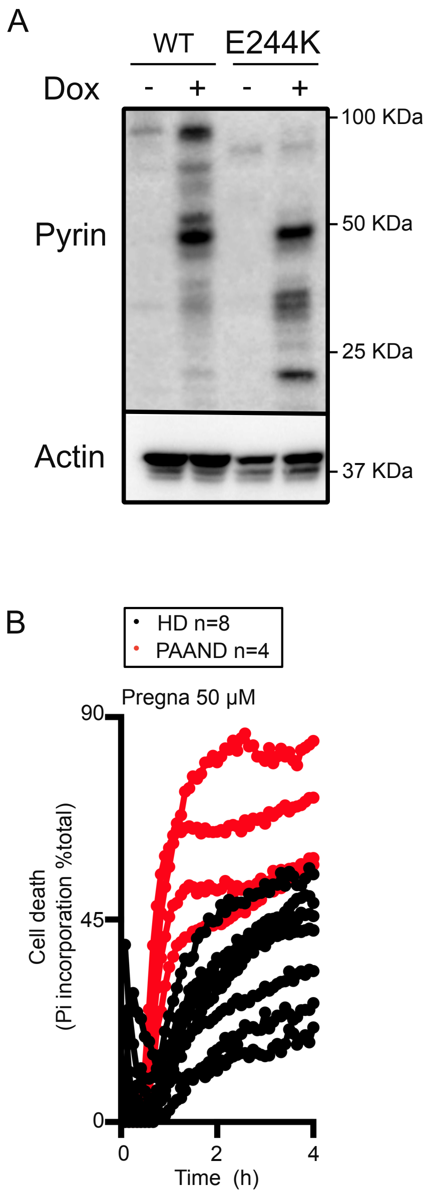
A
WT
E244K
Dox
-
+
-
+
Pyrin
Actin
100 KDa
50 KDa
25 KDa
37 KDa
B
HD n=8
PAAND n=4
Pregna 50 µM
90
45
0
Cell death
(Pi incorporation %total)
0
2
4
Time (h)

Fig. S6

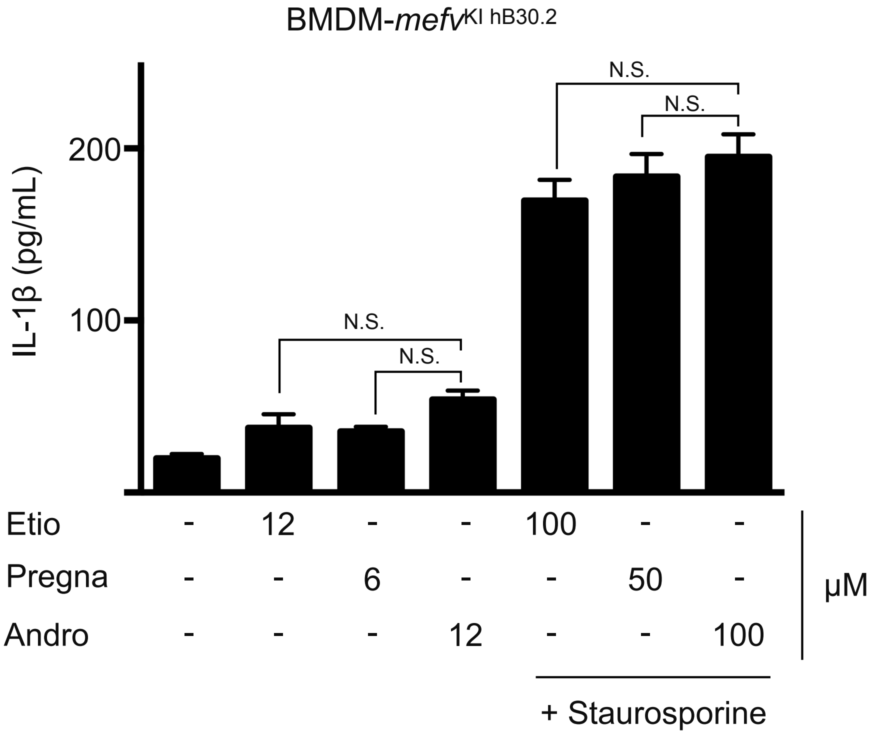
BMDM-mefv KI hB30.2
IL-1β (pg/mL)
200
100
N.S.
N.S.
N.S.
N.S.
Etio - 12 - - 100 - -
Pregna - - 6 - - 50 -
Andro - - - 12 - - 100
μM
+ Staurosporine

Fig. S7

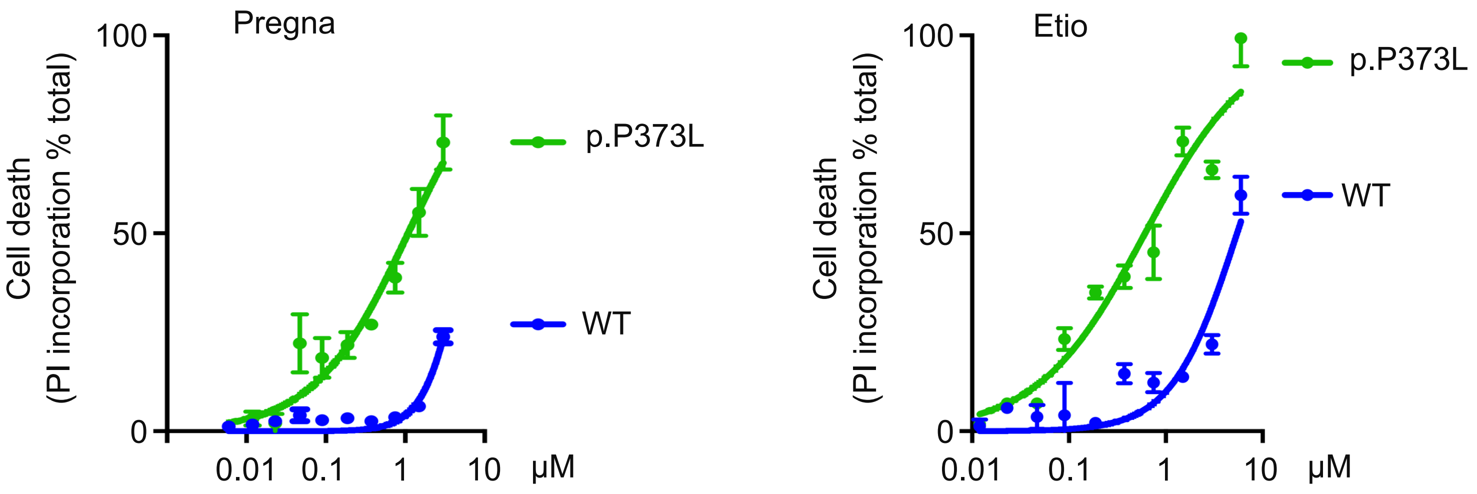
Pregna
100
50
0
Cell death
(PI incorporation % total)
0.01
0.1
1
10
µM
p.P373L
WT
Etio
100
50
0
Cell death
(PI incorporation % total)
0.01
0.1
1
10
µM
p.P373L
WT

Fig. S8

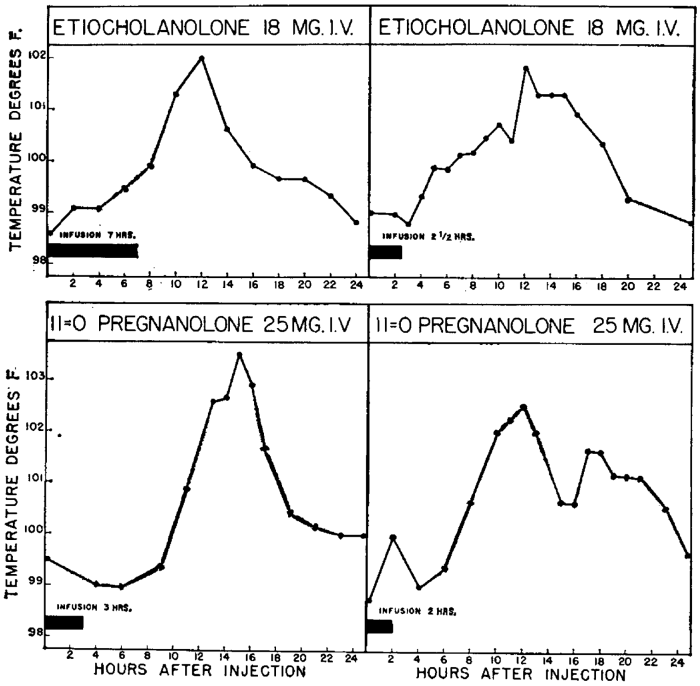
ETIOCHOLANOLONE 18 MG. I.V.
ETIOCHOLANOLONE 18 MG. I.V.
11=O PREGNANOLONE 25 MG. I.V
11=O PREGNANOLONE 25 MG. I.V.
TEMPERATURE DEGREES F.
INFUSION 7 HRS.
INFUSION 2 1/2 HRS.
INFUSION 3 HRS.
INFUSION 2 HRS.
HOURS AFTER INJECTION
HOURS AFTER INJECTION

Fig.S9
